# Integrative transcriptomic profiling of the phosphatome identifies DUSP15 as a chromophobe renal cell carcinoma–selective epithelial biomarker associated with an immune-metabolic tumour state

**DOI:** 10.1101/2025.11.15.688610

**Authors:** Adil R. Sarhan

## Abstract

Chromophobe renal cell carcinoma (chRCC) lacks robust molecular biomarkers and actionable therapeutic targets. Here, 265 protein-coding phosphatases were systematically interrogated using pan-cancer transcriptomic profiles from The Cancer Genome Atlas, 10 independent Gene Expression Omnibus cohorts, pathway and immune analyses, and two independent single-cell RNA-sequencing datasets. *DUSP15* emerged as the most chRCC-selective phosphatase (Tau = 0.977) and distinguished chRCC with high diagnostic accuracy (area under the curve = 0.975; odds ratio = 4.83; P = 6.2 × 10⁻⁷). Cross-cohort validation further reinforced its marked enrichment in chRCC. Transcriptome-wide analyses defined *DUSP15*-high tumours by coordinated enrichment of oxidative phosphorylation, MYC target and DNA repair programs, coupled to attenuation of interferon and inflammatory signalling. Immune deconvolution revealed reduced regulatory T-cell and dendritic-cell signatures and lower overall immune content, identifying an immune-depleted tumour state. Crucially, single-cell analyses resolved the cellular source of the bulk-tumour signal, demonstrating broad *DUSP15* expression throughout the chRCC tumour epithelium in two independent datasets (72.2–83.7% cellular detection; 144.9–204.3 counts per million), with substantially lower expression in immune, stromal, normal renal-epithelial and non-chRCC tumour compartments. Collectively, these findings identify *DUSP15* as a highly selective tumour-epithelial biomarker candidate for chRCC and define a *DUSP15*-associated immune-metabolic state with potential diagnostic and therapeutic relevance.

## Introduction

Chromophobe renal cell carcinoma (chRCC; KICH) is a rare histological subtype of renal cell carcinoma (RCC), accounting for approximately 5% of renal malignancies [1]. Although most localised chRCC tumours follow a relatively indolent clinical course, patients with advanced or metastatic disease have limited therapeutic options and may respond poorly to treatments developed predominantly for clear-cell RCC [1, 2]. Compared with clear-cell and papillary RCC, the molecular determinants of chRCC remain poorly characterised, and validated biomarkers for diagnosis, prognostic stratification and therapeutic selection are lacking [3]. Identifying tumour-selective molecular features of chRCC is therefore important both for improving disease classification and for uncovering biological vulnerabilities that could inform precision therapeutic strategies.

Protein phosphatases are central regulators of intracellular signalling, cellular metabolism, stress responses and immune function [4, 5]. Dual-specificity phosphatases (DUSPs) constitute a functionally diverse phosphatase family capable of dephosphorylating both tyrosine and serine/threonine residues and thereby regulating mitogen-activated protein kinase signalling and other phosphorylation-dependent pathways [6–9]. Several DUSPs have been implicated in tumour growth, treatment resistance and immune regulation [10]; however, the cancer-type specificity and biomarker potential of most members remain incompletely defined. *DUSP15* is among the least characterised members of this family [11], and its cellular distribution, tumour specificity and biological associations in renal cancer have not been systematically established. The distinctive mitochondrial and metabolic biology of chRCC suggests that phosphatase dysregulation could be linked to both tumour-cell state and immune-microenvironment organisation [12]. chRCC tumours frequently exhibit extensive mitochondrial alterations and metabolic reprogramming, yet the relationship between these features and immune exclusion remains unclear [13].

Bulk-tumour transcriptomic profiles can identify disease-selective expression patterns but cannot determine whether a signal originates from neoplastic epithelium, normal renal lineages or infiltrating immune and stromal cells [14]. Resolving this cellular origin is especially important in chRCC because the tumour epithelium retains transcriptional features related to the intercalated-cell lineage, including acid–base transport and mitochondrial programs [15, 16]. Publicly available single-cell RNA-sequencing datasets spanning chRCC, other RCC subtypes and normal kidney provide an opportunity to localise candidate biomarkers at cellular resolution and determine whether their expression is reproduced across independent specimens [17, 18].

Here, 265 protein-coding phosphatases were systematically interrogated using pan-cancer transcriptomic data from The Cancer Genome Atlas [19], independent validation using GEO renal tumour cohorts [20], transcriptome-wide pathway and network analyses, immune-cell deconvolution and two independent renal cancer single-cell RNA-sequencing datasets [17, 18]. *DUSP15* emerged as the most chRCC-selective phosphatase and demonstrated strong diagnostic discrimination between KICH and other renal tumour subtypes. Elevated *DUSP15* expression marked a coordinated oxidative phosphorylation, MYC-target and DNA-repair programme accompanied by reduced inflammatory and immune-cell signatures.

Single-cell analyses independently localised *DUSP15* predominantly to the chRCC tumour epithelium and showed broad expression across neoplastic epithelial cells in two independent datasets. Donor-resolved pathway analyses further revealed a concordant chRCC-versus-ccRCC epithelial pattern characterised by enrichment of oxidative phosphorylation, MYC-target and DNA-repair programmes, together with reduced interferon-γ, TNF-α/NF-κB and inflammatory signalling. Together, these findings position *DUSP15* as a highly selective tumour-epithelial biomarker candidate for chRCC and establish the cellular context of its associated immune–metabolic state, with potential diagnostic and biological relevance.

## Materials and Methods

### Bulk transcriptomic data sources and analysis

A total of 265 protein-coding phosphatase genes were retained for downstream analysis based on manually curated literature-derived lists and Gene Ontology annotations. TCGA gene-expression and clinical data for chromophobe renal cell carcinoma (KICH), kidney renal clear cell carcinoma (KIRC), and kidney renal papillary cell carcinoma (KIRP) were obtained through the UCSC Xena browser and the TCGAbiolinks R package [19]. FPKM expression data retrieved from UCSC Xena were used for renal-subtype expression comparisons and principal component analysis, whereas count-based data retrieved using TCGAbiolinks were used for tumour–normal comparisons, transcriptome-wide correlation analysis, pathway analysis and immune-cell deconvolution.

For independent validation, *DUSP15* expression data were compiled from 10 publicly available Gene Expression Omnibus (GEO) microarray series containing KICH, KIRC, and KIRP samples (GSE2748, GSE11024, GSE11151, GSE12090, GSE16441, GSE17895, GSE19982, GSE36895, GSE46699, and GSE53757) [20]. even datasets contributed 253 renal tumour samples with explicit KICH, KIRC or KIRP annotations and were included in the subtype-specific analyses. The remaining three datasets (GSE36895, GSE46699 and GSE53757) contained normal kidney and RCC samples without reliable histological subtype annotation and were therefore excluded from the KICH–KIRC–KIRP comparisons. Datasets were retrieved using the GEOquery R package. The studies used Affymetrix or Agilent microarray platforms. For each dataset, probe-to-gene mapping was performed using the corresponding platform annotation file, and probes matching the DUSP15 gene symbol were retained. Expression matrices were inspected to determine the reported scale; datasets containing raw or linear-scale expression values were log₂-transformed using *log*₂(*x* + 1), whereas already log₂-normalised datasets were retained on their original scale. Among the 253 subtype-annotated samples, differences in DUSP15 expression across KICH, KIRC and KIRP were assessed using the Kruskal–Wallis test, followed by pairwise Wilcoxon rank-sum tests with Benjamini–Hochberg correction for multiple comparisons.

### Single-cell RNA-sequencing data sources and analysis

Two independent renal cancer single-cell RNA-sequencing datasets were analysed to define the cellular localisation of DUSP15 and assess the reproducibility of its expression in chromophobe renal cell carcinoma (chRCC). GSE152938 was designated as the primary single-cell dataset because it included a chRCC tumour with patient-matched normal kidney together with papillary RCC (pRCC) and clear-cell RCC (ccRCC) specimens [17]. GSE159115 was analysed separately as an independent validation dataset containing chRCC, normal-kidney and ccRCC reference atlases [18]. No cross-study integration or batch correction was performed. Individual cells were used to resolve expression within specimens, whereas donors were treated as the biological units for comparisons between specimens.

For GSE152938, processed 10x Genomics feature-barcode matrices were obtained for GSM4630027–GSM4630031. Specimens were filtered independently using sample-specific detected-gene and mitochondrial-transcript thresholds detailed in Supplementary Information. After quality control, 30,853 cells were retained: 10,133 from the pRCC tumour, 5,667 and 7,251 from the two ccRCC tumours, 7,216 from the chRCC tumour and 586 from matched normal kidney. Each specimen was processed separately in Seurat using library-size normalization to 10,000 counts per cell, log transformation, selection of 2,000 highly variable genes, data scaling, principal-component analysis, shared nearest-neighbour graph construction and Louvain clustering. Sample-specific principal-component and clustering-resolution parameters are provided in Supplementary Information. UMAP was used for annotation review and visualization but not as quantitative evidence.

Clusters were assigned using canonical epithelial, renal-lineage, immune and stromal markers together with the annotation framework of the original study. In the chRCC specimen, clusters C0–C3 comprised 6,454 tumour epithelial cells expressing EPCAM, KRT7, RHCG, FOXI1 and ATP6V1B1. C4–C6 were classified as T/NK cells, tumour-associated macrophages and monocytes, respectively. C7 contained 69 cells with mixed-lineage expression and was excluded from primary compartment comparisons. In matched normal kidney, clusters 0–2 were combined as normal renal epithelium. Tumour epithelial populations in the pRCC and ccRCC specimens were identified using epithelial keratins and subtype-associated markers.

For GSE159115, processed 10x Genomics HDF5 count matrices and deposited cell-annotation tables were obtained from GEO. The single chRCC specimen was GSM4819732. Deposited cell identities were reproduced without de novo reclustering, and per-cell UMI count, detected-gene count and mitochondrial fraction were recalculated for verification. The retained dataset contained 29,474 annotated cells from seven ccRCC donors, one chRCC donor and five normal-kidney donors. The chRCC specimen comprised 2,580 cells, including 2,271 annotated tumour cells. Deposited normal type-A intercalated cells were used as the principal normal epithelial reference, and deposited ccRCC tumour cells provided the non-chRCC tumour reference.

DUSP15 abundance was quantified directly from raw UMI counts. CPM was calculated as the summed DUSP15 UMI count divided by the corresponding total UMI count and multiplied by (10^6). Cellular detection prevalence was defined as the proportion of cells containing at least one DUSP15 UMI. Donor-level expression was calculated as:

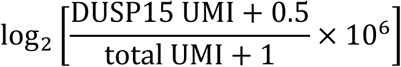

Within GSE152938, the chRCC tumour-epithelial pseudobulk was compared descriptively with patient-matched normal renal epithelium, pRCC tumour epithelium and the equal-donor mean of the two ccRCC tumour-epithelial profiles. Within GSE159115, the chRCC tumour pseudobulk was compared with the equal-donor means of normal type-A intercalated cells from five donors and ccRCC tumour cells from seven donors. Robustness of the chRCC-versus-normal-IC-A contrast was evaluated by sequential normal-donor omission, 1,000 donor-balanced downsampling iterations, restriction to common UMI-depth support and depth-adjusted models. Genome-wide limma–voom analysis was used to position DUSP15 within the distribution of descriptive gene-level effects. Because GSE159115 contained one chRCC donor, these analyses were interpreted as reference-context and gene-ranking assessments rather than population-level hypothesis tests.

To evaluate the immune–metabolic phenotype identified in bulk KICH transcriptomes, seven Hallmark gene sets from MSigDB release 2025.1 were prespecified: oxidative phosphorylation, MYC targets V1, MYC targets V2, DNA repair, interferon-γ response, TNF-α/NF-κB signalling and inflammatory response. Raw counts were aggregated by donor and epithelial compartment. For each gene, the chRCC log₂ pseudobulk value was contrasted with the mean of the indicated reference donors. Competitive pathway enrichment was evaluated using cameraPR with ranked gene-level statistics, and Benjamini–Hochberg false-discovery rates were calculated across the seven pathways. Cross-study concordance was assessed by Spearman correlation of pathway rank-enrichment scores between GSE152938 and GSE159115.

For cell-resolved visualization, counts were normalized to 10,000 UMIs per cell and log-transformed. Each pathway score was calculated as the mean normalized expression of its detected member genes and residualized for log-transformed total UMI and detected-gene counts. Standardized oxidative-phosphorylation, MYC-target and DNA-repair scores were averaged to generate a metabolic/proliferative composite, whereas interferon-γ, TNF-α/NF-κB and inflammatory-response scores were averaged to generate an immune/inflammatory composite. Continuous, depth-adjusted associations between DUSP15 expression and the composite scores were summarized using Spearman correlation without defining an arbitrary DUSP15-positive threshold. For UMAP visualization, colour limits were restricted to the 2nd and 98th percentiles; untruncated values were used in all quantitative summaries. Cell-level associations were interpreted descriptively because cells from the same specimen are not independent biological replicates.

### Tissue Specificity Analysis

Tissue specificity for each gene in the datasets was quantified using the Tau index, a measurement of selective expression across tissues [21]. The Tau score was calculated using the equation:

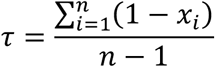

Where *x_i_* represents the expression in tissue *i*, normalized to the gene maximum expression. Whereas *n* represents the number of tissues. Tau values ranging from 0 (ubiquitous expression) to 1 (perfect specificity).

### Differential Expression Analysis

Differential expression of *DUSP15* between KICH tumours and adjacent normal tissues in the TCGA-KICH dataset was assessed using both Wilcoxon rank-sum tests (log₂-transformed FPKM) and DESeq2-normalized counts. Genes with FDR-adjusted p < 0.05 and log₂ fold change > 1.5 were considered significant.

### Diagnostic Performance and Logistic Regression

To evaluate diagnostic potential of *DUSP15* and other DUSP genes, Receiver Operating Characteristic (ROC) analysis in the TCGA and GEO microarray datasets were performed using the pROC package in R. Binary classification was defined as KICH = 1 and non-KICH (KIRC + KIRP) = 0. Area Under the Curve (AUC) values were generated, with additional ROC analysis comparing TCGA-KICH vs. normal tissues (tumour = 1, normal = 0). For *DUSP15*, binary logistic regression was applied using log₂-transformed values to estimate odds ratios (OR), 95% confidence intervals (CI), and p-values.

### Principal Component Analysis

PCA was conducted on expression values of DUSP genes across KICH, KIRC, and KIRP tumours using TCGA cohort. Expression matrices were scaled and centred using the prcomp function in R. PCA plots were annotated by cancer subtype, and 90% confidence ellipses were overlaid. Loadings and explained variance for each principal component were reported.

### Clinical and Pathologic Correlation

*DUSP15* expression was analysed across AJCC tumour stage, pathological T/N/M categories, histological variants, tumour laterality, patient age, and survival status using the TCGA-KICH dataset. Statistical comparisons were made using Kruskal-Wallis tests and Wilcoxon rank-sum tests. Post hoc analyses were adjusted using the Benjamini-Hochberg method. Tumour volume was estimated using the ellipsoid formula [22]:

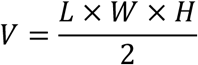

Where *V* is the tumour volume in cubic centimetres (cm³). *L*, *W*, and *H*represent the tumour’s maximum length, width, and height, respectively. Only samples with complete dimensional data were included. Spearman correlation was used to assess associations between tumour volume, age at diagnosis and *DUSP15* expression.

### Transcriptome-Wide Correlation and Gene Set Enrichment Analysis

Spearman correlation was computed between *DUSP15* and all protein-coding genes across TCGA-KICH tumours. Genes were ranked by correlation coefficients, and those with |*ρ*| > 0.3 and FDR < 0.05 were considered significant. Gene Set Enrichment Analysis (GSEA) [23] was performed using Hallmark gene sets from MSigDB (v2025.1.Hs) via the clusterProfiler R package. Enriched pathways (FDR < 0.05) were used to construct Gene-Concept Networks by linking core enriched genes to their respective terms. Networks were visualized in Cytoscape (v3.10.3) [24] using the yFiles Organic Layout, with pathway nodes colour-mapped by normalized enrichment score (NES).

### Immune Deconvolution

The xCell algorithm was applied to TCGA-KICH tumours to estimate relative enrichment of 64 immune and stromal cell types [25]. Tumours were grouped into DUSP15-Low (Q1) and DUSP15-High (Q4) categories. ImmuneScore and MicroenvironmentScore were compared between groups using Wilcoxon tests. FDR correction was applied to all comparisons.

### Software and Visualization

Analyses and graphical outputs were generated primarily in R. The final single-cell RNA-sequencing analyses were performed in R version 4.6.0 using data.table 1.18.4, Matrix 1.7-5, Seurat 5.5.0, SeuratObject 5.4.0, edgeR 4.10.0, limma 3.68.2 and rhdf5 2.56.0. TCGAbiolinks and GEOquery were used to retrieve TCGA and GEO data, respectively. Differential expression analysis of bulk transcriptomic data was performed using DESeq2. Receiver operating characteristic curves and corresponding areas under the curve were calculated using pROC. Gene set enrichment analysis was conducted using clusterProfiler with gene annotations obtained from org.Hs.eg.db, and immune-cell deconvolution was performed using xCell.

**Table 1:** ROC AUC and mean expression of DUSP family genes in KICH (TCGA Cohort).

| DUSP Gene | AUC | Mean Exp. in KICH |
| --- | --- | --- |
| DUSP15 | 0.9756 | 51.417 |
| DUSP11 | 0.9301 | 2.8419 |
| DUSP8 | 0.9270 | 11.768 |
| DUSP12 | 0.8939 | 3.3643 |
| DUSP19 | 0.8920 | 0.5876 |
| DUSP14 | 0.8875 | 4.6169 |
| DUSP22 | 0.8803 | 3.8270 |
| DUSP18 | 0.8761 | 0.6643 |
| DUSP23 | 0.8645 | 30.489 |
| DUSP5 | 0.8180 | 6.3210 |
| DUSP10 | 0.8118 | 1.8465 |
| DUSP1 | 0.7770 | 80.133 |
| DUSP16 | 0.7679 | 12.430 |
| DUSP26 | 0.7132 | 0.4907 |
| DUSP7 | 0.6988 | 2.4002 |
| DUSP9 | 0.6862 | 4.4429 |
| DUSP3 | 0.6838 | 26.501 |
| DUSP2 | 0.6487 | 2.5054 |
| DUSP4 | 0.6247 | 3.8607 |
| DUSP28 | 0.6210 | 0.9567 |
| DUSP6 | 0.6067 | 15.375 |
| DUSP13B | 0.5730 | 0.0208 |
| DUSP21 | 0.4804 | 0.0008 |
| DUSP29 | 0.4414 | 0.0076 |

Statistical graphics were generated using ggplot2 4.0.3 and ggpubr, and heatmaps were produced using pheatmap. Random seeds were fixed before single-cell clustering, UMAP generation and repeated downsampling to support reproducibility. Gene–concept networks were constructed from the core-enrichment genes identified by gene set enrichment analysis. Network data were processed in Python version 3.11 using pandas and NetworkX and subsequently visualised in Cytoscape version 3.10.3. The yFiles Organic Layout was used to organise network topology and improve visual clarity.

## Results

### DUSP15 defines a highly specific transcriptomic signature of chromophobe renal cell carcinoma among human phosphatases

A systematic pan-cancer analysis was conducted to identify protein phosphatases with tumour-type selective expression across 21 cancer types from TCGA. Tissue specificity was quantified using the Tau score, a stringent metric for transcriptomic restriction (Supplementary Table S1). This unbiased screen revealed a discrete subset of phosphatases exhibiting near-exclusive expression in single tumour types, including *ACP3* (Tau = 0.999), a known prostate cancer marker [26], and *G6PC1* (Tau = 0.998), recently reported as a liver-specific biomarker in hepatocellular carcinoma [27]. Among dual-specificity phosphatases, *DUSP15* ranked highest with a Tau score of 0.977, showing strong and selective expression in chromophobe renal cell carcinoma. Heatmap visualization of z-score-normalized expression for the top 50 most tissue-specific phosphatases confirmed the unique enrichment of *DUSP15* in KICH compared to all other tumour types (Fig. 1A).

**Fig 1.**
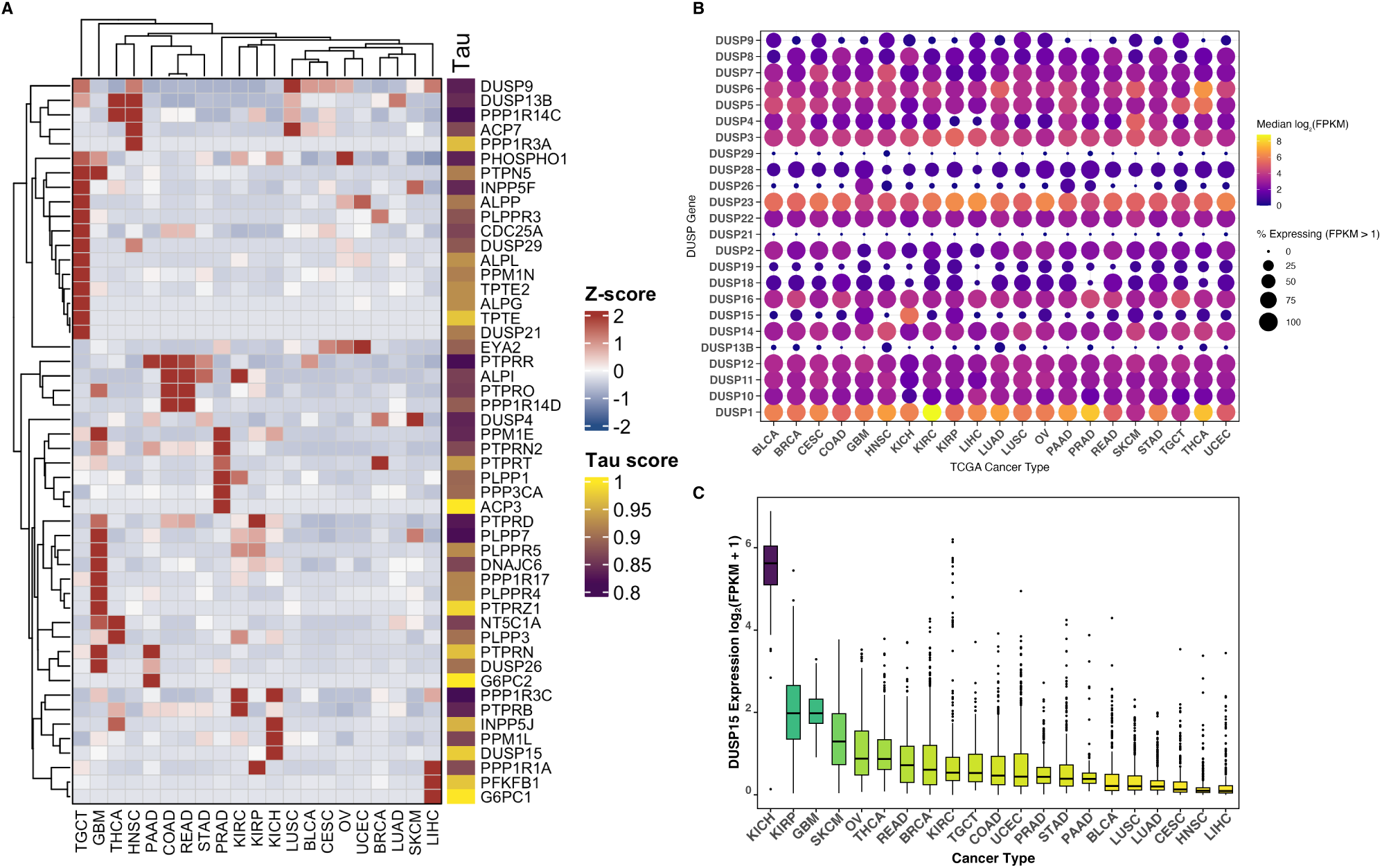
DUSP15 is a tumour-selective phosphatase with exceptional specificity for chromophobe renal cell carcinoma. (A) Heatmap of z-score normalized expression for the top 50 most tissue-specific protein phosphatases across 21 tumour types in TCGA. Genes were ranked by Tau score, a continuous metric of tissue specificity (0-1), with higher values indicating more restricted expression. DUSP15 emerged as the most KICH-selective dual-specificity phosphatase (Tau = 0.977), showing strong expression in KICH and minimal expression elsewhere. (B) Dot plot of median expression (log₂ FPKM) and proportion of expressing samples (% with FPKM > 1) for all DUSP family members across TCGA tumours. DUSP15 was detected in 98.4% of KICH samples and was notably absent or low in most other cancers. (C) Boxplot of DUSP15 expression across 21 TCGA cancer types. Kruskal-Wallis testing indicated significant inter-tumour variation (P < 0.0001). Dunn’s post hoc comparisons (Benjamini-Hochberg adjusted) confirmed significantly elevated expression in KICH compared to multiple cancer types, including LIHC, LUAD, COAD, and HNSC (P values < 10⁻³¹). See Supplementary Table S2 for full statistics. Cancer types are denoted using TCGA abbreviations as follows: BLCA, bladder urothelial carcinoma; BRCA, breast invasive carcinoma; CESC, cervical squamous cell carcinoma and endocervical adenocarcinoma; COAD, colon adenocarcinoma; GBM, glioblastoma multiforme; HNSC, head and neck squamous cell carcinoma; KICH, kidney chromophobe; KIRC, kidney renal clear cell carcinoma; KIRP, kidney renal papillary cell carcinoma; LIHC, liver hepatocellular carcinoma; LUAD, lung adenocarcinoma; LUSC, lung squamous cell carcinoma; OV, ovarian serous cystadenocarcinoma; PAAD, pancreatic adenocarcinoma; PRAD, prostate adenocarcinoma; READ, rectum adenocarcinoma; SKCM, skin cutaneous melanoma; STAD, stomach adenocarcinoma; TGCT, testicular germ cell tumours; THCA, thyroid carcinoma; and UCEC, uterine corpus endometrial carcinoma.

To further contextualize this selectivity, the expression of the entire DUSP family was examined across the TCGA cohort. Among 21 tumour types, *DUSP15* was uniquely and robustly upregulated in KICH, with a median expression of 5.62 log₂ (FPKM) and detection in 98.4% of KICH samples (Fig. 1B). In contrast, most other DUSPs (e.g., *DUSP1, DUSP3, DUSP5*) were broadly expressed across multiple malignancies with low tissue specificity. These results identify *DUSP15* as a molecular outlier among DUSPs, reinforcing its potential as a tumour-selective biomarker.

Quantitative statistical analysis confirmed the strength of this tumour specificity. Kruskal-Wallis testing revealed highly significant variation in *DUSP15* expression across cancers (P < 0.0001). Pairwise comparisons using Dunn’s test with Benjamini-Hochberg correction demonstrated that *DUSP15* expression in KICH was significantly higher than in nearly all other tumour types, including liver hepatocellular carcinoma (LIHC; adjusted P = 4.85 × 10⁻⁸⁴), lung adenocarcinoma (LUAD; P = 1.43 × 10⁻⁶⁵), and colon adenocarcinoma (COAD; P = 3.06 × 10⁻³¹) (Fig. 1C; Supplementary Table S2). The most pronounced difference was observed with head and neck squamous cell carcinoma (HNSC) (Z = –20.83; P = 2.34 × 10⁻⁹⁵). Collectively, these results define *DUSP15* as one of the most tumour-restricted phosphatases in the TCGA transcriptome, with exceptional specificity for chromophobe renal cell carcinoma.

### DUSP15 is a KICH-specific dual-specificity phosphatase with superior discriminative power among renal tumours

To further delineate the tissue and subtype specificity of *DUSP15*, its expression was systematically analysed across the three major histological subtypes of renal carcinoma using TCGA RNA-seq datasets. *DUSP15* exhibited striking overexpression in KICH compared to both clear cell renal cell carcinoma (KIRC) and papillary renal cell carcinoma (KIRP) (Fig. 2A). A Kruskal-Wallis test revealed a highly significant difference in *DUSP15* expression across the subtypes (χ² = 588.74, df = 2, *P* < 2.2 × 10⁻¹⁶). Post hoc pairwise Wilcoxon tests with Benjamini-Hochberg correction confirmed that *DUSP15* levels were significantly higher in KICH compared to KIRC (adj. P = 3.06 × 10⁻³⁵) and KIRP (adj. P = 8.97 × 10⁻³³), and also significantly different between KIRP and KIRC (adj. P = 2.64 × 10⁻⁶⁶). These results establish *DUSP15* as a highly selective phosphatase marker for KICH, distinctively enriched relative to other kidney cancer subtypes.

**Fig 2.**
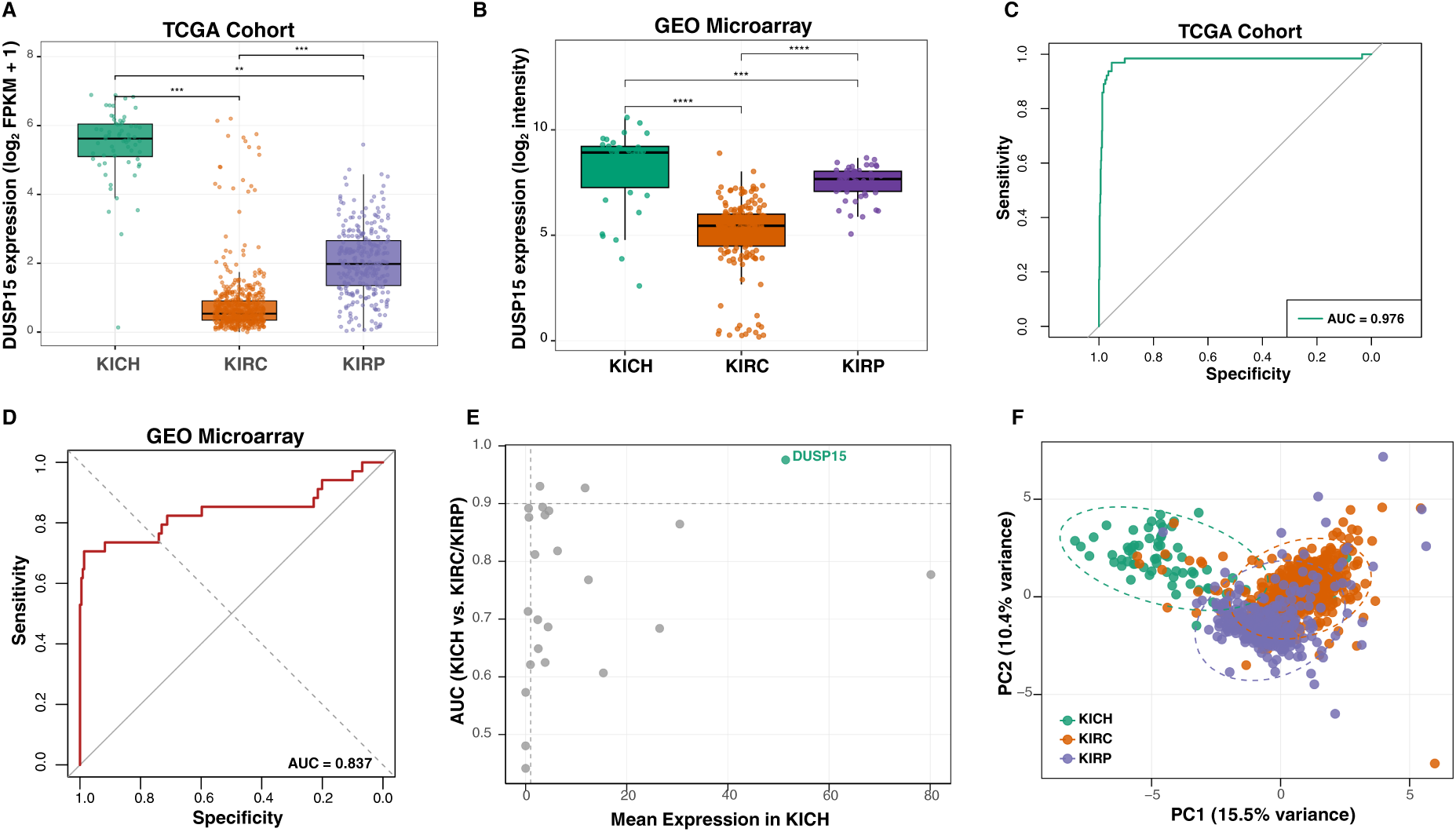
DUSP15 is a KICH-specific dual-specificity phosphatase with exceptional diagnostic power. (A) DUSP15 expression across TCGA renal carcinoma subtypes, including KIRC, KIRP and KICH, shows pronounced upregulation in KICH. Kruskal-Wallis testing confirmed significant differences across subtypes (χ² = 588.74, df = 2, P < 2.2 × 10⁻¹⁶). Post hoc pairwise Wilcoxon tests with Benjamini-Hochberg correction showed DUSP15 was significantly elevated in KICH compared to KIRC (P = 3.06 × 10⁻³⁵) and KIRP (P = 8.97 × 10⁻³³), with additional differences between KIRC and KIRP (P = 2.64 × 10⁻⁶⁶). (B) Independent validation using 253 subtype-annotated renal tumour samples from seven GEO datasets confirmed enrichment of DUSP15 in KICH. Expression was highest in KICH (mean = 8.15), intermediate in KIRP (7.50), and lowest in KIRC (5.08). Kruskal–Wallis test: P = 5.87 × 10⁻²⁶. Benjamini–Hochberg-adjusted pairwise Wilcoxon P values were 2.24 × 10⁻¹¹ for KICH versus KIRC and 1.54 × 10⁻⁴ for KICH versus KIRP. (C) Receiver operating characteristic (ROC) analysis comparing the performance of DUSP15 (AUC = 0.976) in distinguishing KICH from KIRC and KIRP using TCGA cohort. (D) ROC analysis in the GEO datasets yielded an AUC of 0.837 (95% CI: 0.735–0.939; P = 1.06 × 10⁻¹⁰), validating DUSP15’s diagnostic potential. Optimal threshold: 8.45; sensitivity = 70.6%, specificity = 98.6%. (E) Scatterplot of mean FPKM expression in TCGA-KICH versus AUC for each DUSP gene. (F) PCA based on DUSP gene expression revealed clear separation of KICH along PC1 (15.5% variance explained), with DUSP15 contributing the highest negative loading (–0.394), underscoring its role in subtype-specific transcriptomic divergence.

To validate the TCGA findings independently, *DUSP15* expression was evaluated in 253 renal tumour samples with explicit KICH, KIRC or KIRP annotations drawn from seven GEO microarray datasets within the 10-dataset compilation (Supplementary Table S3). *DUSP15* expression was highest in KICH (mean ± SD = 8.15 ± 1.93), intermediate in KIRP (7.50 ± 0.77), and lowest in KIRC (5.08 ± 1.72) (Fig. 2B). A Kruskal–Wallis test confirmed significant overall differences among the three subtypes (H = 116.20, df = 2, P = 5.87 × 10⁻²⁶). Pairwise Wilcoxon rank-sum tests with Benjamini–Hochberg correction showed significant differences between KICH and KIRC (adjusted P = 2.24 × 10⁻¹¹), KICH and KIRP (adjusted P = 1.54 × 10⁻⁴), and KIRP and KIRC (adjusted P = 1.43 × 10⁻²¹).

To benchmark *DUSP15* against other members of the dual-specificity phosphatase family, its diagnostic performance was evaluated using Receiver operating characteristic (ROC) analysis across all detected DUSP genes in TCGA-KICH, -KIRC, and -KIRP tumours (Table 1). Among all family members, *DUSP15* achieved the highest area under the curve (AUC = 0.976), indicating exceptional diagnostic accuracy for distinguishing KICH from non-KICH renal tumours (Fig. 2C). By comparison, *DUSP11* (AUC = 0.930), *DUSP8* (AUC = 0.927), and *DUSP12* (AUC = 0.894) also showed promising performance, but none matched the sensitivity and specificity of *DUSP15*. Most other DUSPs exhibited modest to poor diagnostic capacity, with several, including *DUSP21* and *DUSP29*, falling below the performance threshold (AUC < 0.50). These data establish *DUSP15* as the most reliable and discriminative biomarker for KICH within the DUSP family. To validate the ROC findings from the TCGA cohort in an independent dataset, the same classification analysis was performed for *DUSP15* using the integrated GEO kidney cancer dataset. ROC curve analysis yielded an AUC of 0.837 (95% CI: 0.735-0.939; p = 1.06 × 10⁻¹⁰) for distinguishing KICH from other subtypes (Fig 2D). The optimal expression threshold of 8.45 provided a sensitivity of 70.6% (95% CI: 52.5-84.9%) and a specificity of 98.6% (95% CI: 96.0-99.7%). These findings independently validate the selective upregulation and diagnostic power of *DUSP15* in KICH observed in the TCGA cohort, reinforcing its potential as a robust biomarker for chromophobe renal cell carcinoma.

To evaluate the correlation between expression abundance and diagnostic accuracy, mean FPKM expression in TCGA-KICH was plotted against the AUC for each DUSP gene (Fig. 2E). *DUSP15* uniquely occupied the high-expression, high-AUC quadrant, underscoring its ideal combination of detectability and specificity. Other high-AUC candidates such as *DUSP11* and *DUSP8* resided in the low-expression range, which may limit their translational utility. This analysis further reinforces *DUSP15* as a uniquely advantageous biomarker, one that couples robust expression with nearly perfect classification potential. To explore whether *DUSP15* also contributes to transcriptional divergence between renal cancer subtypes, principal component analysis (PCA) was performed using the expression profiles of the DUSP genes across KICH, KIRC, and KIRP using TCGA cohort (Supplementary Table S4). PCA revealed clear segregation of KICH tumours from the other two subtypes along the first principal component (PC1), which explained 15.5% of total variance (Fig. 2F). *DUSP15* exhibited the strongest negative loading on PC1 (–0.394), indicating that it was the dominant contributor to this axis of variation. While *DUSP11* and *DUSP12* also contributed meaningfully, their influence was comparatively lower. Taken together, these results establish *DUSP15* not only as a potent diagnostic biomarker for chromophobe RCC but also as a key molecular driver of KICH-specific gene expression programs within the phosphatase landscape.

### DUSP15 expression is maintained across clinical and demographic strata but modestly declines in advanced KICH

To evaluate whether *DUSP15* expression is modulated by clinicopathologic or demographic variables, its distribution was analysed across TCGA-KICH clinical annotations. Consistent with its diagnostic utility, *DUSP15* expression remained robustly elevated across most subgroups. Stratification by AJCC pathological stage revealed a modest but statistically significant reduction in stage IV tumours compared to earlier stages (Fig. 3A), suggesting potential transcriptional remodelling in a subset of advanced cases. Analysis by pathological T stage demonstrated a significant decrease in *DUSP15* expression between T2b and T4 tumours (adjusted P = 0.0418), and a near-significant reduction between T2 and T1a (adjusted P = 0.0511) (Fig. 3C), while other comparisons showed no significant changes, indicating overall stability across early stages. Although reduced *DUSP15* levels were observed in N1 and M1 tumours relative to N0 and M0 groups, the extremely limited sample sizes (n = 2) preclude definitive interpretation (Fig. 3D, 3E). Expression levels in NX and MX cases were comparable to early-stage tumours, further suggesting that declines in N1/M1 reflect sample variability rather than consistent trends.

**Fig 3.**
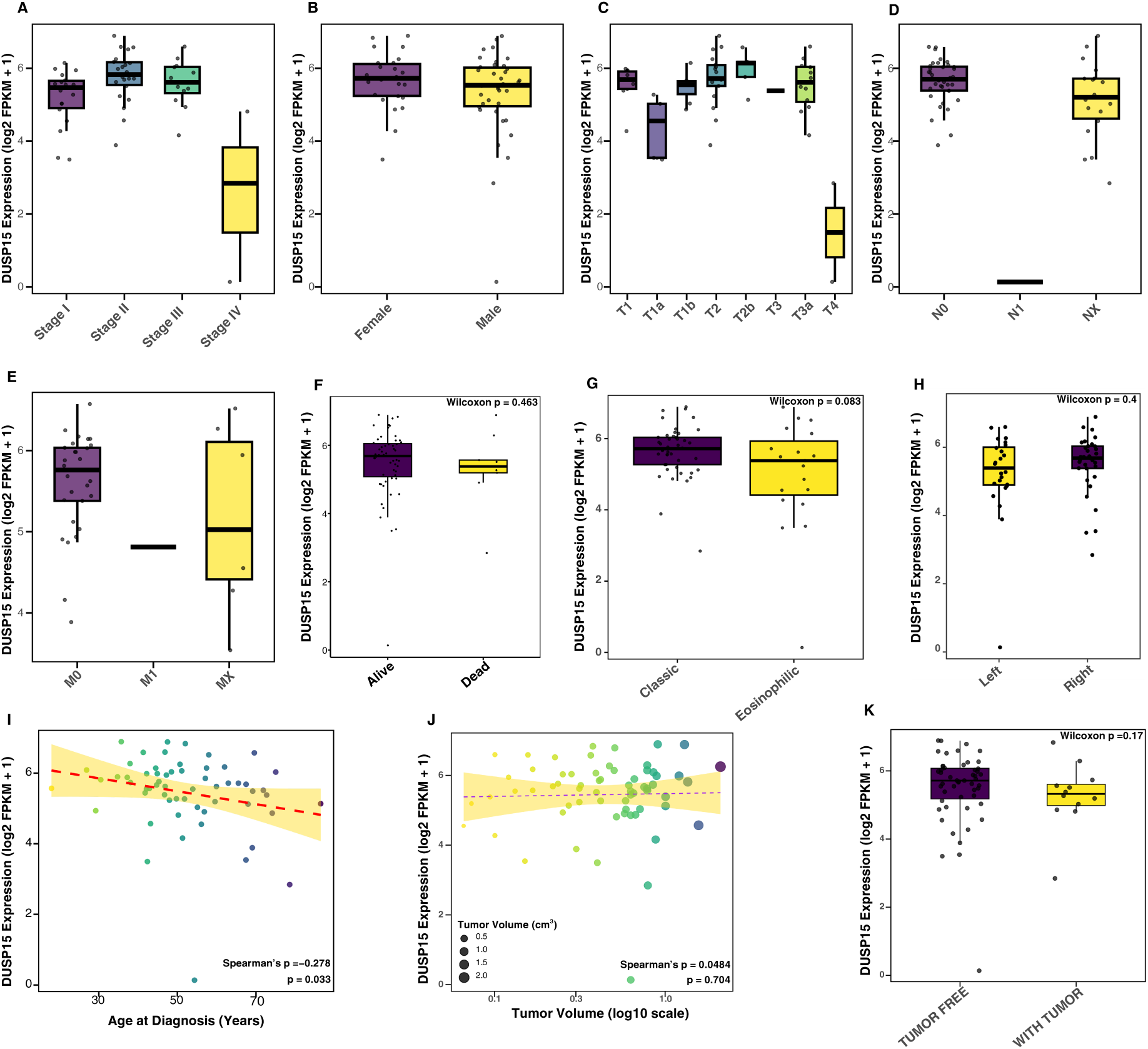
Stability of DUSP15 expression across KICH clinical subgroups. (A) DUSP15 expression across AJCC pathological stages in TCGA-KICH. A modest but significant reduction is observed in stage IV tumours compared to early stages. (B) DUSP15 expression stratified by patient gender; no significant difference was detected. (C) Comparison of DUSP15 expression across pathological T stages; significant difference observed between T2b and T4 tumours. (D, E) Expression of DUSP15 stratified by nodal (N) and metastatic (M) stages shows a dip in N1 and M1. (F) DUSP15 expression in patients alive versus deceased at last follow-up; no significant difference observed. (G) Expression comparison between eosinophilic and classic KICH histological subtypes; no statistically significant difference. (H) DUSP15 levels in tumours located in the left versus right kidney show no significant laterality bias. (I) Spearman correlation of DUSP15 expression with age at diagnosis reveals a modest but significant negative correlation. (J) Correlation between DUSP15 expression and tumour volume (cm³). (K) Expression levels in tumour-free versus tumour-present cases at data collection timepoint. All statistical tests were two-sided, with multiple comparisons corrected using the Benjamini-Hochberg method where applicable.

Demographic variables had minimal impact: no significant sex-based differences were observed (P = 0.91; Fig. 3B), and only a weak inverse correlation was detected with age at diagnosis (Spearman ρ = –0.278, P = 0.033; Fig. 3I). *DUSP15* expression was comparable between classic and eosinophilic histological variants (P = 0.083; Fig. 3G) and unaffected by tumour laterality (P = 0.396; Fig. 3H). Furthermore, no differences were detected with respect to patient survival status (P = 0.463; Fig. 3F), disease activity at sampling (P = 0.17; Fig. 3K), or estimated tumour volume (ρ = 0.048, P = 0.704; Fig. 3J). Collectively, these results demonstrate that *DUSP15* expression is largely stable across clinical and demographic contexts in KICH, with only minor reductions observed in advanced-stage disease and older patients. Its consistent expression across histological subtypes, anatomical locations, and tumour burden reinforces its utility as a robust, qualitative diagnostic marker for chromophobe renal cell carcinoma.

### DUSP15 is a quantitatively predictive and tumour-selective diagnostic biomarker in KICH

To evaluate the diagnostic potential of *DUSP15* in TCGA-KICH, its expression levels between primary tumour tissues and adjacent normal tissue was compared (Supplementary Table S5; Fig. 4A). This analysis revealed a highly significant upregulation of *DUSP15* in tumour tissues, as assessed by a Wilcoxon rank-sum test with continuity correction (P = 3.81 × 10⁻¹²), supporting its robust differential expression and selective enrichment in KICH tumours. To assess the capacity of *DUSP15* to discriminate tumour from normal tissue, ROC curve analysis was conducted (Fig. 4B). The resulting AUC was 0.973, indicating near-perfect sensitivity and specificity. This exceptional classification performance confirms *DUSP15*’s strong discriminatory power and further supports its role as a diagnostic biomarker in chromophobe renal tumours.

**Fig 4.**
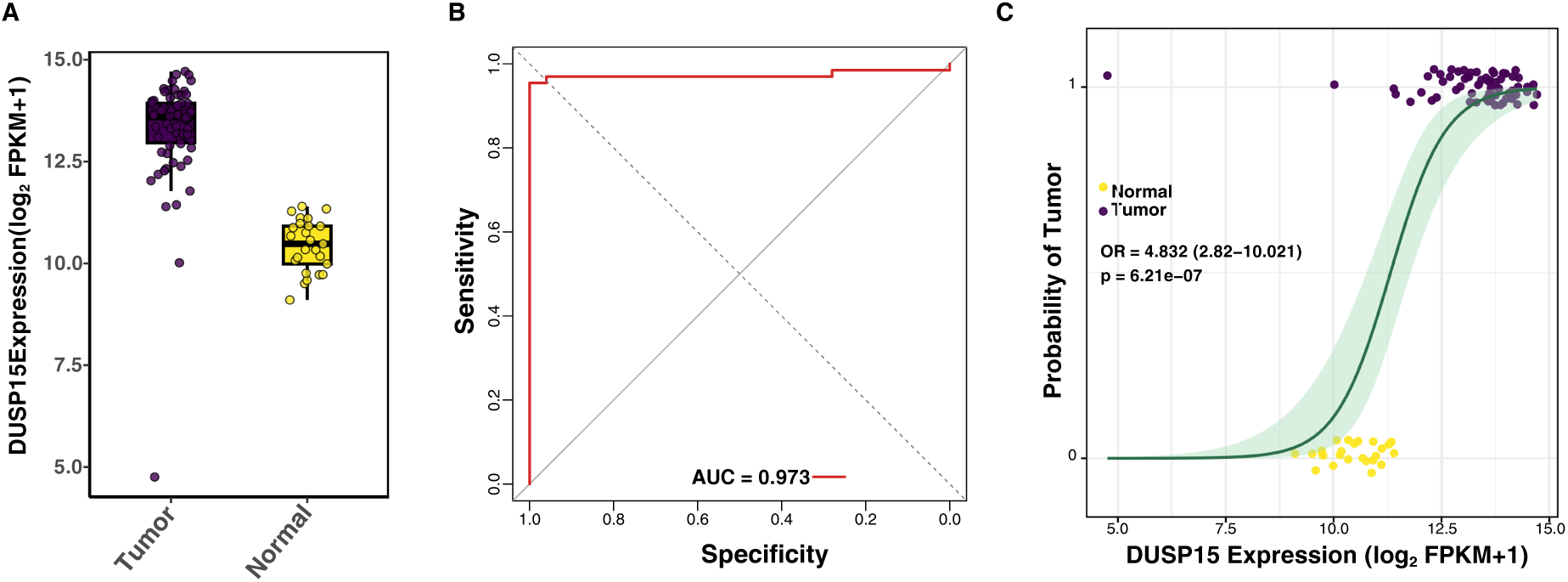
DUSP15 is a robust diagnostic biomarker in KICH. (A) Expression of DUSP15 in primary chromophobe renal cell carcinoma (KICH) tumours versus adjacent normal tissue from TCGA revealed a marked and significant upregulation in tumours. Wilcoxon rank-sum test with continuity correction: P = 3.81 × 10⁻¹² (Supplementary Table S5). (B) ROC curve analysis demonstrated exceptional diagnostic performance of DUSP15 for distinguishing tumour from normal tissue, with an AUC of 0.973. (C) Binary logistic regression analysis using log₂-transformed expression of DUSP15 showed a strong association with tumour status, with each unit increase in expression corresponding to a 4.83-fold increase in the odds of KICH diagnosis (95% CI: 2.82–10.02; P = 6.21 × 10⁻⁷).

To quantify the strength of this diagnostic association, a binary logistic regression model was employed using log₂-transformed *DUSP15* expression levels as the predictor variable (Fig. 4C). The model revealed that each one-unit increase in *DUSP15* expression was associated with a 4.83-fold increase in the odds of KICH diagnosis (Table 2).

**Table 2:**
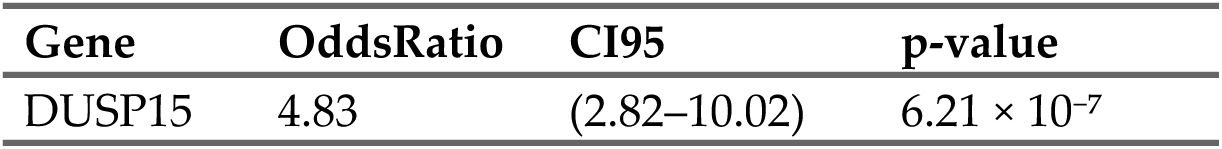
Logistic regression showing odds ratio, CI, and significance comparing KICH vs. normal tissue (TCGA Cohort).

| Gene | OddsRatio | CI95 | p-value |
| --- | --- | --- | --- |
| *DUSP15* | 4.83 | (2.82–10.02) | $6.21 \times 10^{-7}$ |

These data establish that *DUSP15* expression is not only significantly elevated in KICH tumours but also quantitatively predictive of tumour status, with excellent diagnostic precision. Together, these findings position *DUSP15* as a highly specific and statistically robust biomarker for kidney chromophobe carcinoma, with significant potential for integration into diagnostic frameworks aimed at distinguishing malignant from normal renal tissue.

### DUSP15 expression defines a metabolic-immune dichotomy in KICH

To delineate the molecular landscape associated with *DUSP15* expression in KICH, a genome-wide Spearman correlation analysis was conducted using RNA-seq data from TCGA-KICH tumours. This analysis identified 1,881 genes positively correlated and 2,194 genes negatively correlated with *DUSP15* expression, indicating that *DUSP15* is associated with broad transcriptional variation across TCGA-KICH tumours (Supplementary Table S6).

To systematically characterize *DUSP15*-associated transcriptional programs, Gene Set Enrichment Analysis (GSEA) was performed using Hallmark gene sets, ranking all genes by their Spearman correlation with *DUSP15* expression (Supplementary Table S7). The enrichment profile of *DUSP15*-high tumours was dominated by mitochondrial and biosynthetic programs, with oxidative phosphorylation emerging as the top-enriched pathway (NES = 2.881, FDR = 4.17 × 10⁻¹⁰), alongside upregulation of MYC targets, cholesterol homeostasis, and DNA repair. Conversely, *DUSP15* expression was negatively associated with multiple immune and inflammatory signatures, including interferon gamma response, TNF-α signalling via NF-κB, and KRAS signalling. To visualize the functional connectivity of these pathways, a Gene-Concept Network was constructed by linking significant GSEA terms (FDR < 0.05) to their core enriched genes. The resulting bipartite network revealed two distinct modules (Fig. 5): a tightly clustered metabolic arm composed of positively enriched pathways, and a large, interconnected immune-suppression module dominated by negatively enriched signatures. This structural separation underscores the transcriptional duality of *DUSP15*, which appears to promote metabolic activation while concurrently repressing pro-inflammatory signalling in KICH.

**Fig 5.**
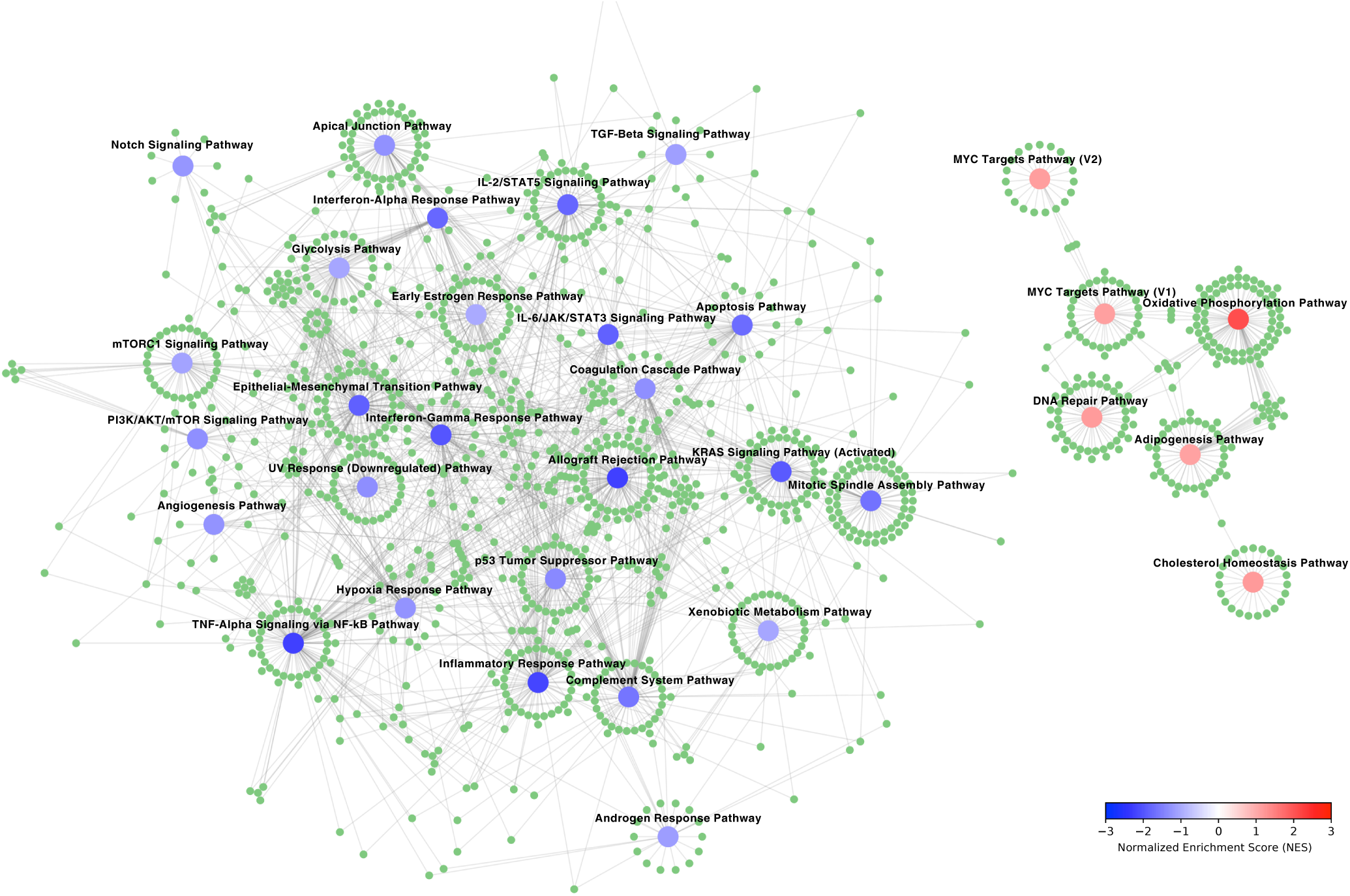
Gene-Concept Network of significantly enriched Hallmark pathways, illustrating the transcriptional landscape associated with DUSP15 expression. The network was visualized in Cytoscape using the yFiles Organic Layout, which optimizes spatial organization based on node connectivity to minimize edge overlap and highlight functional modules. Pathway nodes are colour-coded by normalized enrichment score (NES), and gene nodes are uniformly coloured.

### Transcriptional co-regulation and immune depletion characterize DUSP15-high tumours

To complement pathway-level insights from GSEA and the gene-concept network, *DUSP15*-associated gene expression programs were interrogated at single-gene resolution using genome-wide Spearman correlation analysis in TCGA-KICH tumours. The 50 most positively and 50 most negatively correlated protein-coding genes were extracted, z-score normalized, and visualized across *DUSP15* expression quartiles, revealing marked expression stratification between *DUSP15*-high (Q4) and *DUSP15*-low (Q1) groups (Fig. 6). Among the top positively correlated genes, *SPTBN4* (*ρ* = 0.744, adjusted p = 2.94 × 10⁻⁹), a cytoskeletal adaptor, showed the strongest association with *DUSP15* (Fig. 6A). Mitochondrial and biosynthetic factors such as MRPL34, *MICOS13*, *MRPL54*, and *NDUFB7* (*ρ* ≈ 0.69–0.71; adjusted p < 10⁻⁷) were prominently enriched, suggesting enhanced translation, cristae organization, and metabolic activation in *DUSP15*-high tumours (Fig. 6A). Additional co-expressed genes included PPDPF, a Ras/MAPK pathway modulator, and the apoptosis regulator *FEM1A*, as well as splicing components (*SNRNP25*, *YJU2*) and the long non-coding RNA *LINC03073*, highlighting a tightly co-regulated biosynthetic axis.

**Fig 6.**
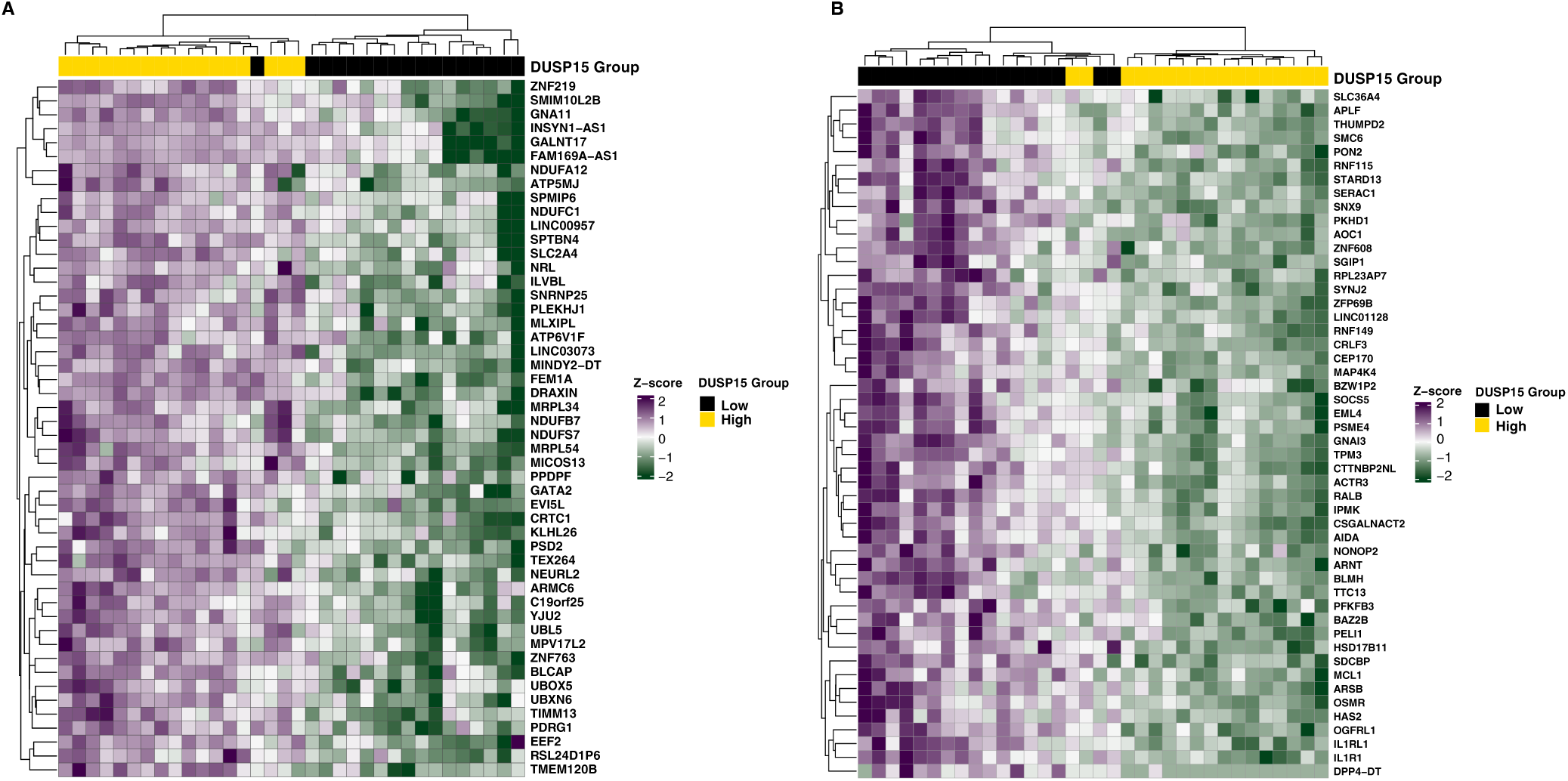
Biosynthetic and immune-silent programs associated with DUSP15 in KICH. (A) Heatmap showing z-score–normalized expression of the top 50 positively correlated genes with DUSP15 across TCGA-KICH tumour samples. (B) Heatmap showing z-score–normalized expression of the top 50 negatively correlated genes with DUSP15 across TCGA-KICH tumour samples. Samples were grouped by DUSP15 expression quartiles (Q1 = low, Q4 = high). Hierarchical clustering highlights distinct expression profiles between DUSP15-High and DUSP15-Low groups.

In contrast, negatively correlated genes were enriched for immune and cytokine signalling programs (Fig. 6B). *IL1R1* (*ρ* = –0.673) and its decoy receptor *IL1RL1*/ST2 (*ρ* = –0.652) were among the most repressed, alongside *SGIP1* (endocytosis), *AOC1* (immune recruitment), and *HSD17B11* (oxidative stress), all with *ρ* > 0.63 and adjusted p < 2 × 10⁻⁵. Additional suppressed transcripts included *SYNJ2* (phosphoinositide signalling), the DNA repair factor APLF (*ρ* = –0.619), and STARD13, a Rho-GTPase–activating protein, collectively suggesting attenuation of inflammatory sensing, stress responses, and cytoskeletal remodelling.

To determine whether this transcriptional signature was mirrored by changes in the tumour microenvironment, xCell deconvolution was applied to TCGA-KICH transcriptomic data (Supplementary Table S8; Fig. 7). Tumours were stratified by *DUSP15* quartiles, and enrichment scores were compared between Q4 and Q1. *DUSP15*-high tumours exhibited a markedly immune-depleted phenotype. The global ImmuneScore was significantly reduced (median = 0.0079 vs. 0.0254; P = 1.3 × 10⁻⁴, FDR = 0.0034), and the MicroenvironmentScore was similarly attenuated (median = 0.0255 vs. 0.0739; P = 8.7 × 10⁻⁴, FDR = 0.0116), indicating broad suppression of immune and stromal components. Specific immune populations such as conventional dendritic cells (cDCs) and regulatory T cells (Tregs) were significantly depleted in *DUSP15*-high tumours, consistent with an immune-excluded microenvironment.

**Fig 7.**
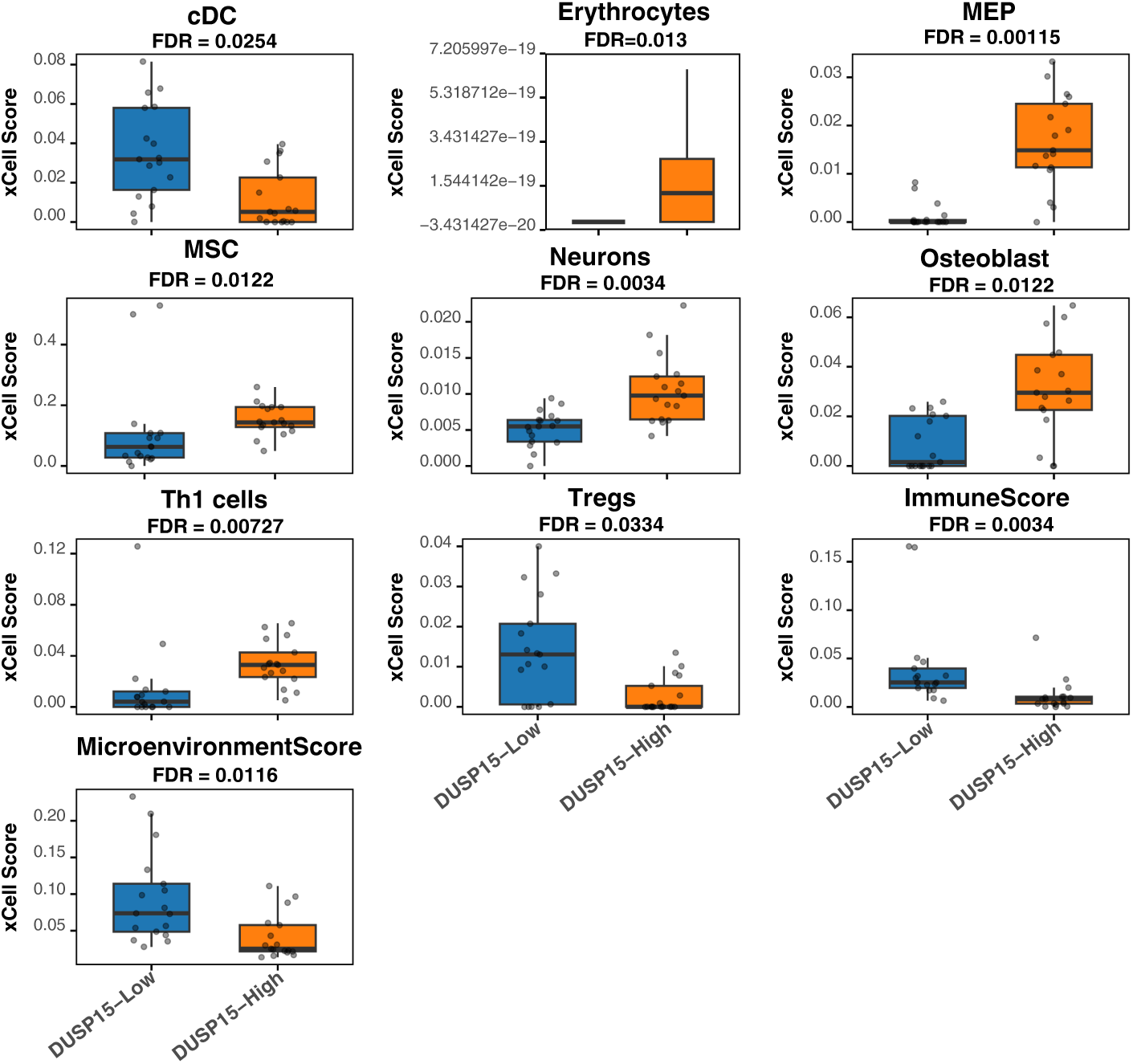
Immune and stromal remodelling in DUSP15-high chromophobe renal tumours. xCell deconvolution analysis of TCGA-KICH transcriptomes revealed that tumours with high DUSP15 expression (Q4) exhibit a profoundly immune-depleted phenotype relative to low-expressing tumours (Q1). The ImmuneScore (median = 0.0079 vs. 0.0254; P = 1.3 × 10⁻⁴, FDR = 0.0034) and MicroenvironmentScore (median = 0.0255 vs. 0.0739; P = 8.7 × 10⁻⁴, FDR = 0.0116) were both significantly reduced. Key immune populations such as conventional dendritic cells and regulatory T cells were markedly suppressed. In contrast, stromal and progenitor signatures, including megakaryocyte-erythroid progenitors (FDR = 0.0012), mesenchymal stem cells (FDR = 0.0122), and erythrocytes (FDR = 0.0130), were enriched in DUSP15-high tumours. Boxplots display median and interquartile ranges, with statistical significance assessed using Wilcoxon rank-sum tests and FDR-adjusted p-values indicated.

Unexpectedly, *DUSP15*-high tumours demonstrated enrichment of non-immune stromal lineages, including megakaryocyte-erythroid progenitors (MEPs), neuronal signatures, and mesenchymal stem cells (MSCs). MEPs showed a striking increase (P = 1.7 × 10⁻⁵, FDR = 0.0012), and MSCs were similarly elevated (P = 1.1 × 10⁻³, FDR = 0.0122), suggesting stromal reprogramming and potential erythroid skewing. Erythrocyte signatures were also higher in *DUSP15*-high tumours (P = 1.6 × 10⁻³, FDR = 0.0130), further supporting this hypothesis. While Th1 cells were unexpectedly enriched (FDR = 0.0073), most immune subsets were depleted, reinforcing the presence of a noncanonical immune-desert state.

Together, these findings define a dual-axis program associated with high *DUSP15* expression in KICH: intrinsic transcriptional upregulation of mitochondrial and biosynthetic machinery, and extrinsic remodelling of the tumour microenvironment toward immune exclusion and stromal transformation. This immune-metabolic reprogramming highlights *DUSP15* as a potential regulator of tumour adaptation and a target for therapeutic intervention.

### Single-cell RNA-seq analyses localise DUSP15 to the chRCC tumour epithelium and define its immune–metabolic context

To validate the tumour specificity of DUSP15 observed in bulk transcriptomic analyses and resolve its cellular distribution within the tumour microenvironment, two independent single-cell RNA-seq datasets were analysed. Analysis of Single-cell RNA-seq dataset (GSE152938) retained 30,853 cells across five independently processed specimens. The chRCC tumour contributed 7,216 cells, of which 6,454 formed four marker-supported tumour-epithelial clusters (Supplementary Fig. S1 and Supplementary Tables S9–S10). These clusters co-expressed epithelial and intercalated-cell-lineage markers, including *EPCAM*, *KRT7*, *RHCG*, *FOXI1* and *ATP6V1B1*, and were separated from *PTPRC*-, LYZ- and *NKG7*-expressing immune populations (Fig. 8A,C). *DUSP15* expression mapped predominantly to the tumour-epithelial region of the UMAP (Fig. 8B) and was distributed across all four tumour-epithelial clusters rather than being restricted to one transcriptional cluster. Cluster-level abundance ranged from 100.6 to 165.0 CPM, with *DUSP15* detected in 54.0–82.2% of tumour epithelial cells (Fig. 8D; Supplementary Fig. S3C and Supplementary Table S11).

**Figure 8.**
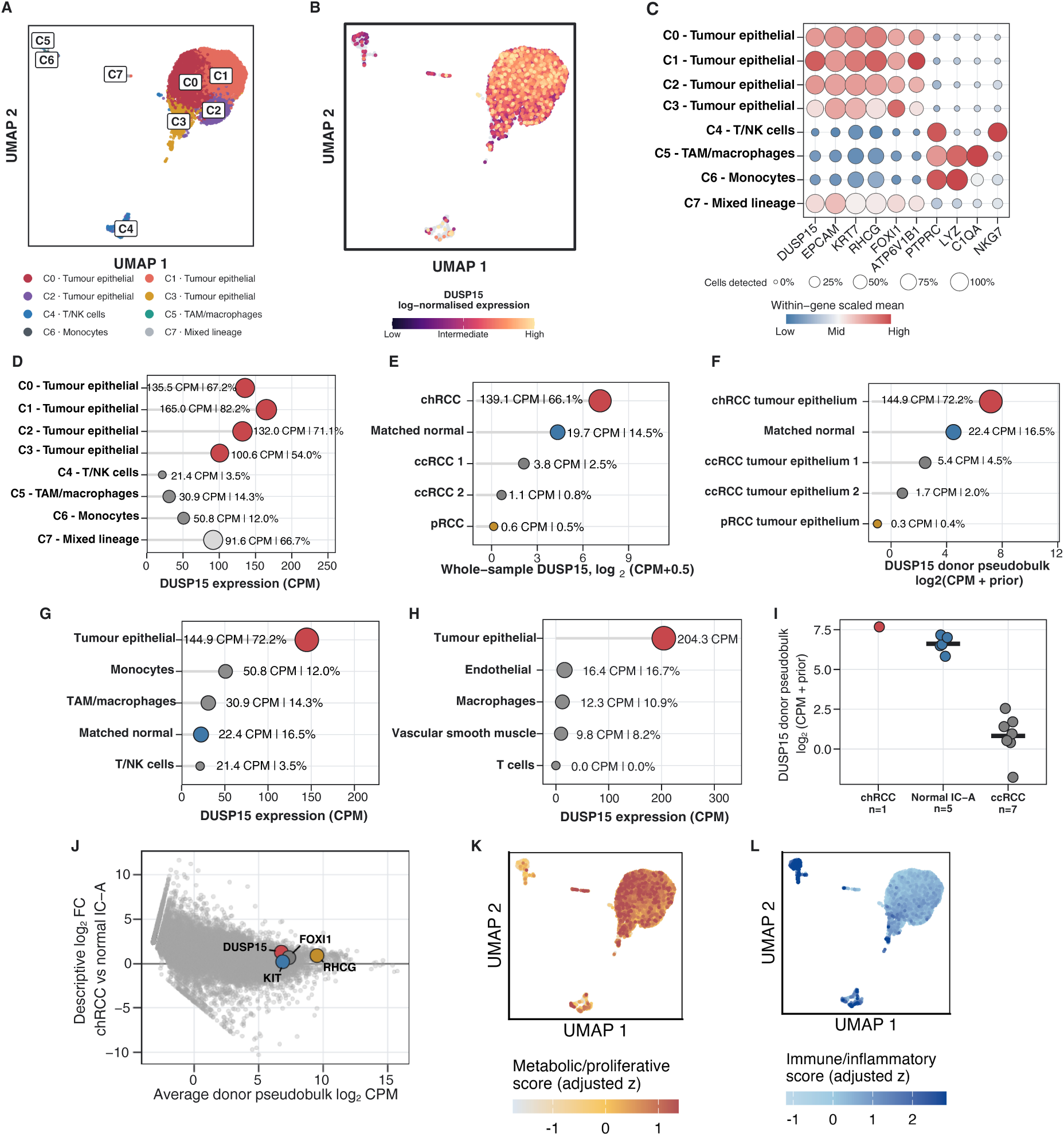
**Single-cell localization and immune–metabolic context of DUSP15 in chromophobe renal cell carcinoma. (**A) UMAP of 7,216 cells from the GSE152938 chRCC specimen, coloured by marker-supported cluster identity. C7 was retained for visualization but excluded from the primary quantitative comparisons. (B) DUSP15 expression projected onto the same UMAP embedding; colour denotes log-normalized expression. (C) Marker dot plot supporting the cluster assignments. Point area represents the percentage of cells detecting each transcript, and colour represents mean log-normalized expression scaled independently within each gene. (D) DUSP15 abundance and detection prevalence across the eight GSE152938 chRCC clusters. (E) Whole-sample DUSP15 expression across the five GSE152938 specimens, comprising one chRCC tumour, its matched normal kidney, two ccRCC tumours and one pRCC tumour. Horizontal position represents log₂(CPM + 0.5); labels report untransformed CPM and the percentage of cells with at least one DUSP15 UMI. (F) Tumour-epithelial DUSP15 pseudobulk expression in the GSE152938 chRCC specimen, matched normal renal epithelium, two ccRCC specimens and one pRCC specimen. (G) DUSP15 abundance across the principal cellular compartments of the GSE152938 chRCC specimen, with matched normal renal epithelium included for comparison. (H) DUSP15 abundance across annotated cellular compartments in the independent GSE159115 chRCC specimen; four unassigned cells were omitted. (I) Donor-level DUSP15 pseudobulk expression in GSE159115 chRCC tumour epithelium (n = 1 donor), normal type-A intercalated cells (IC-A; n = 5 donors) and ccRCC tumour epithelium (n = 7 donors). Each point represents one donor; horizontal bars denote reference-group means. Log₂ CPM values were calculated using a 0.5-UMI prior count. (J) Genome-wide descriptive relationship between average donor-pseudobulk expression and the chRCC-versus-normal-IC-A log₂ fold change in GSE159115. DUSP15 and the lineage-associated genes KIT, RHCG and FOXI1 are highlighted for biological context. (K, L) GSE152938 chRCC UMAPs coloured by depth-adjusted metabolic/proliferative (K) and immune/inflammatory (L) composite scores.

At the whole-sample level, the chRCC specimen showed the highest *DUSP15* abundance in the GSE152938 cohort (139.1 CPM; 66.1% cellular detection), compared with matched normal kidney (19.7 CPM; 14.5%), two ccRCC specimens (3.8 and 1.1 CPM) and the pRCC specimen (0.6 CPM) (Fig. 8E). Restriction to tumour epithelial cells strengthened this disease-context pattern. The chRCC tumour epithelium contained 10,404 *DUSP15* UMIs, corresponding to 144.9 CPM, with expression detected in 72.2% of 6,454 cells. In contrast, *DUSP15* abundance was 22.4 CPM in matched normal renal epithelium, 5.4 and 1.7 CPM in the two ccRCC tumour-epithelial populations and 0.34 CPM in the pRCC tumour-epithelial population (Fig. 8F; Supplementary Table S12). On the donor-pseudobulk scale, the chRCC value was 6.43-fold higher than matched normal renal epithelium, 46.7-fold higher than the mean of the two ccRCC donors and 286.8-fold higher than the pRCC specimen (Supplementary Table S12).

Compartment-level analysis localised the increased *DUSP15* signal to the chRCC tumour epithelium. *DUSP15* abundance and cellular detection were substantially lower in monocytes (50.8 CPM; 12.0% detected), tumour-associated macrophages (30.9 CPM; 14.3%), T/NK cells (21.4 CPM; 3.5%) and patient-matched normal renal epithelium (22.4 CPM; 16.5%) than in chRCC tumour epithelial cells (144.9 CPM; 72.2%) (Fig. 8G; Supplementary Table S11). The concordant UMAP, marker and raw-count pseudobulk results therefore identify *DUSP15* as a broadly expressed tumour-epithelial transcript within this chRCC specimen.

The independent GSE159115 dataset reproduced this localisation pattern. Among 2,580 cells from the chRCC specimen, the deposited tumour-epithelial compartment comprised 2,271 cells and contained 6,762 *DUSP15* UMIs. This corresponded to 204.3 CPM, with *DUSP15* detected in 83.7% of tumour cells. Expression was markedly lower in endothelial cells (16.4 CPM; 16.7% detected), macrophages (12.3 CPM; 10.9%), vascular smooth-muscle cells (9.8 CPM; 8.2%) and T cells (0 CPM; 0%) (Fig. 8H; Supplementary Fig. S3D and Supplementary Tables S13–S15). Together, the two independent datasets showed closely convergent tumour-epithelial *DUSP15* profiles (Supplementary Table S18).

Donor-level reference comparisons further established the expression context of the GSE159115 chRCC specimen. *DUSP15* abundance was 2.09-fold higher than the equal-donor mean of normal IC-A cells from five donors and 115.4-fold higher than the equal-donor mean of ccRCC tumour cells from seven donors (Fig. 8I; Supplementary Table S17). The positive chRCC-versus-normal-IC-A contrast was retained after omission of each normal donor (log₂ fold-change range, 0.87–1.20), donor-balanced down sampling (median log₂ fold change, 1.06; 2.5th–97.5th percentile interval, 0.60–1.56), restriction to common UMI-depth support (log₂ fold change, 0.97) and depth-adjusted modelling (Supplementary Fig. S2A and Supplementary Table S17). Genome-wide contextualisation placed *DUSP15* on the positive side of the chRCC-versus-normal-IC-A effect distribution (Fig. 8J), whereas its chRCC-versus-ccRCC effect ranked at the 99.1st percentile of genome-wide descriptive effects (Supplementary Fig. S2B and Supplementary Table S16).

To determine whether the immune–metabolic phenotype associated with *DUSP15* in bulk KICH transcriptomes was detectable within the epithelial compartment, seven Hallmark pathways were prespecified for donor-resolved pseudobulk analysis (Supplementary Tables S19–S20). Relative to ccRCC tumour epithelium, the chRCC specimen in each dataset showed positive rank enrichment of oxidative phosphorylation, MYC targets V1, MYC targets V2 and DNA repair, accompanied by negative enrichment of interferon-γ response, TNF-α/NF-κB signalling and inflammatory response. All seven pathways exhibited the prespecified direction and reached a false-discovery rate below 0.05 in both index-donor comparisons (Supplementary Tables S21a–S21b). Their enrichment-score ordering was identical between GSE152938 and GSE159115 for the chRCC-versus-ccRCC contrast (Spearman *ρ* = 1.00), showing strong cross-study concordance (Supplementary Fig. S4 and Supplementary Table S22). Relative to normal renal epithelial references, the four metabolic/proliferative pathways also showed positive enrichment in both chRCC specimens. However, the three immune/inflammatory pathways were positively rather than negatively enriched, and cross-study pathway concordance was weaker (Spearman *ρ* = 0.214). Thus, the single-cell data reproduced the bulk immune–metabolic direction most clearly when chRCC tumour epithelium was compared with ccRCC tumour epithelium, rather than demonstrating universal immune suppression relative to normal renal epithelium (Supplementary Fig. S4 and Supplementary Tables S21a–S22). Because each dataset contributed one chRCC donor, these comparisons assess reproducibility between two index specimens and do not estimate a population-level chRCC effect.

Cell-resolved programme scores revealed a corresponding compartmental organisation. In GSE152938, the mean depth-adjusted metabolic/proliferative composite was higher in tumour epithelial cells than in the immune microenvironment (mean adjusted z score, 0.10 versus −1.01), whereas the immune/inflammatory composite showed the inverse pattern (−0.18 versus 1.71) (Fig. 8K,L). The same directional separation was observed in GSE159115, with mean metabolic/proliferative scores of 0.11 in tumour epithelium and −0.86 in microenvironmental cells, and mean immune/inflammatory scores of −0.24 and 1.78, respectively (Supplementary Table S23). Within each chRCC specimen, however, *DUSP15* expression showed only weak cell-level correlations with the composite scores after depth adjustment (absolute Spearman *ρ* ≤ 0.135). These findings establish the compartmental context of *DUSP15* but do not imply that variation in *DUSP15* expression directly determines programme intensity among individual tumour cells (Supplementary Table S24).

## Discussion

Chromophobe renal cell carcinoma (chRCC; KICH) remains a comparatively understudied kidney cancer subtype, with limited molecular biomarkers and few precision therapeutic strategies for patients with advanced disease [28, 29]. By integrating phosphatase-wide expression profiling, independent cross-cohort validation, pathway and immune analyses, and two single-cell RNA-sequencing datasets, this study identifies *DUSP15* as a highly selective and biologically informative chRCC biomarker candidate. *DUSP15* exhibited the greatest chRCC selectivity among the dual-specificity phosphatases examined and distinguished KICH from clear-cell and papillary RCC with an area under the curve of 0.976 in TCGA. Independent analysis of GEO microarray cohorts produced an area under the curve of 0.837, supporting the reproducibility of *DUSP15* enrichment across cohorts and expression platforms. Although these results require prospective clinical validation, they position *DUSP15* as a strong candidate for molecular classification of chRCC.

The biological significance of *DUSP15* extended beyond its diagnostic performance. Transcriptome-wide correlation and gene set enrichment analyses associated elevated *DUSP15* expression with oxidative phosphorylation, MYC target and DNA repair programs. These pathways are consistent with the mitochondrial abundance, metabolic specialisation and stress-adaptation mechanisms characteristic of chRCC. Conversely, *DUSP15* expression was inversely associated with interferon-γ response, TNF-α signalling and other inflammatory programs. Gene–concept network analysis organised the core-enrichment genes into interconnected metabolic and immune-related modules, illustrating the coordinated structure of the *DUSP15*-associated transcriptional state. These findings support an association between *DUSP15* expression, enhanced mitochondrial-metabolic programs and attenuated immune signalling; however, they do not demonstrate that *DUSP15* enzymatic activity directly regulates these pathways.

At the gene level, positive *DUSP15* correlates included mitochondrial ribosomal components, ion-transport genes and RNA-processing factors, linking *DUSP15* expression to biological programs that support bioenergetic adaptation and cellular homeostasis [30–32]. Negative correlates included genes involved in cytokine signalling, stress responses and extracellular-matrix regulation. Consistent with these transcriptional relationships, xCell analysis associated *DUSP15*-high tumours with lower inferred regulatory T-cell and dendritic-cell enrichment and reduced overall immune and microenvironment scores. These results support an immune-depleted phenotype in *DUSP15*-high KICH tumours. Because deconvolution methods estimate relative transcriptional enrichment rather than directly measuring cell abundance, histological and spatial validation will be required to establish the organisation of the *DUSP15*-associated tumour microenvironment.

The single-cell analyses established the cellular context of *DUSP15* expression. In the primary GSE152938 dataset, *DUSP15* was broadly expressed throughout the chRCC tumour-epithelial compartment, with 10,404 *DUSP15* UMIs corresponding to 144.9 counts per million and detection in 72.2% of 6,454 tumour-epithelial cells. Expression extended across all four marker-supported tumour-epithelial clusters and was substantially higher than in matched normal renal epithelium, immune compartments, pRCC tumour epithelium and ccRCC tumour epithelium. The independent GSE159115 dataset reproduced this localisation: its chRCC tumour compartment contained 6,762 *DUSP15* UMIs, corresponding to 204.3 counts per million, with detection in 83.7% of 2,271 tumour cells. Expression was markedly lower in endothelial, macrophage, vascular smooth-muscle and T-cell compartments. Donor-level pseudobulk comparisons further demonstrated enrichment relative to normal type-A intercalated cells and ccRCC tumour cells. The concordance between these independently generated datasets indicates that the elevated bulk-tumour *DUSP15* signal is principally attributable to broad expression within the chRCC neoplastic epithelium rather than to immune or stromal infiltration.

The single-cell pathway analysis further clarified the immune–metabolic transcriptional context associated with *DUSP15* in bulk KICH tumours. Relative to ccRCC tumour epithelium, both datasets reproduced positive enrichment of oxidative phosphorylation, MYC-target and DNA-repair programmes together with negative enrichment of interferon-γ response, TNF-α/NF-κB signalling and inflammatory-response programmes. All seven prespecified pathways were significant in both datasets, and their enrichment scores showed complete rank-order concordance between studies. These findings provide independent cell-resolved support for a metabolically active, immune-attenuated chRCC epithelial state relative to ccRCC. Importantly, the immune component was reference dependent: although metabolic and proliferative programmes remained enriched relative to normal renal epithelium, immune and inflammatory pathways were also positively enriched in that comparison. The single-cell findings therefore support immune attenuation specifically in relation to ccRCC rather than a universal reduction relative to normal renal epithelium. Cell-resolved programme scores also distinguished tumour epithelial cells from immune and stromal compartments, with tumour epithelium showing higher metabolic/proliferative and lower immune/inflammatory scores in both chRCC specimens. Each single-cell study contributed one chRCC donor; accordingly, these data demonstrate reproducible cellular localisation and pathway context across two independent specimens but do not estimate inter-patient variability or population-level single-cell effects. Larger chRCC single-cell cohorts will be required to determine how *DUSP15* expression and its associated transcriptional programmes vary with tumour stage, morphology, genomic alterations and clinical outcome.

Collectively, this study identifies DUSP15 as a highly selective and reproducible chRCC-associated phosphatase that is predominantly localised to tumour-associated epithelial populations. Its expression marks a coordinated tumour state characterised by mitochondrial, MYC-target and DNA-repair enrichment, together with reduced immune and inflammatory signatures relative to ccRCC. These findings support further evaluation of DUSP15 as a candidate molecular marker for distinguishing chRCC from other major renal cancer subtypes and provide a focused rationale for mechanistic investigation of its relationship with tumour metabolism, immune regulation and chRCC epithelial identity.

## Supporting information

Supplementary Information for single cell analysis

## Declarations

### Ethics approval and consent to participate

Not applicable.

## Data Availability

All datasets analysed in this study are publicly available. TCGA bulk RNA-seq expression and clinical data for chromophobe renal cell carcinoma (KICH), kidney clear cell carcinoma (KIRC), and kidney papillary carcinoma (KIRP) were obtained from the UCSC Xena browser (https://xenabrowser.net). Pan-cancer TCGA expression data were retrieved from the Genomic Data Commons (GDC) data portal (https://portal.gdc.cancer.gov).

Gene Expression Omnibus (GEO) microarray datasets used for independent validation were obtained from the National Center for Biotechnology Information (NCBI) GEO repository (https://www.ncbi.nlm.nih.gov/geo/) under accession numbers GSE2748, GSE11024, GSE11151, GSE12090, GSE16441, GSE17895, GSE19982, GSE36895, GSE46699, and GSE53757.

The single-cell RNA-sequencing datasets analysed in this study are publicly available through the Gene Expression Omnibus under accession numbers GSE152938 and GSE159115.

## Competing interests

The author declares no competing interests.

## Funding

This research received no external funding or institutional financial support.

## Authors’ contributions

A.R.S. conceived and designed the study, performed all analyses, interpreted the results, prepared the figures, and wrote the manuscript.

## Acknowledgements

The author gratefully acknowledges the researchers and consortia who generated and made publicly available the datasets used in this study, including The Cancer Genome Atlas (TCGA), the Gene Expression Omnibus (GEO), and the investigators of the GSE152938 and GSE159115 single-cell RNA-sequencing datasets.

## Code availability

The complete analysis code, audited metadata, processed source-data tables, computational environment information, and scripts used to generate the main and supplementary figures are available at https://github.com/adilsarhan/DUSP15-KICH-single-cell. The version associated with this article is archived as release v1.0.0 at https://github.com/adilsarhan/DUSP15-KICH-single-cell/releases/tag/v1.0.0.

## Notes

### Competing Interest Statement

The authors have declared no competing interest.

### Summary of Updates

This version has been revised to improve methodological and interpretive precision. The GEO validation design and the use of different TCGA expression data sources have been clarified, including the sample numbers used in each analysis. The description of the single-cell RNA sequencing results has been refined to distinguish tumour-associated epithelial populations from definitively malignant cells. Interpretation of the immune deconvolution and pathway analyses has also been made more cautious to reflect their transcriptome-derived and associative nature. The exploratory SIGLEC15 correlation analysis and related claims have been removed. The title, abstract, main text, figures and supplementary files have been updated accordingly. These revisions do not alter the central conclusion that DUSP15 is selectively enriched in chromophobe renal cell carcinoma and is predominantly associated with tumour-associated epithelial populations and an immune-metabolic transcriptional state.

