## Supplementary Information for single cell analysis for "Integrative transcriptomic profiling of the phosphatome identifies DUSP15 as a chromophobe renal cell carcinoma–selective epithelial biomarker associated with an immune-metabolic tumour state"

Adil R. Sarhan

Department of Medical Laboratory Techniques, Nasiriyah Technical Institute, Southern Technical University, Nasiriyah 64001, Iraq

#### Overview

This Supplementary Information provides the complete analytical framework supporting the single-cell RNA-sequencing component of the study. GSE152938 was used as the primary dataset because it contains chRCC with patient-matched normal kidney and additional pRCC and ccRCC specimens. GSE159115 was evaluated independently to test whether the tumour-epithelial DUSP15 pattern was reproduced in a separate study. The supplement reports quality control, compartment assignment, raw-count DUSP15 summaries, donor-level reference comparisons, sensitivity analyses and cross-study replication.

### Supplementary Methods

#### Dataset selection and independent-study design

Two publicly available renal cell carcinoma scRNA-seq datasets were analysed separately. GSE152938 [1] provided one pRCC tumour, two ccRCC tumours, one chRCC tumour and patient-matched normal kidney. GSE159115 [2] provided independently generated normal-kidney, ccRCC and chRCC atlases. No cross-study integration or batch correction was performed; reproducibility was assessed by comparing results obtained independently within each study.

#### GSE152938 quality control and processing

Processed 10x Genomics feature-barcode matrices were obtained for GSM4630027–GSM4630031. Cells were retained at 200–5,000 detected genes with mitochondrial transcripts ≤10% for pRCC and ccRCC or ≤30% for chRCC. The matched normal sample was filtered at 200–2,500 genes and mitochondrial transcripts ≤30%. Each specimen was analysed separately in Seurat. Counts were library-size normalised to 10,000 per cell and log-transformed; 2,000 variable features were selected using the variance-stabilising transformation method, scaled and subjected to principal-component analysis. Neighbour graphs and Louvain clusters used the sample-specific parameters reported in Supplementary Table S10. UMAP embeddings were used to review annotation quality but not as quantitative evidence.

#### GSE152938 compartment assignment

Clusters were assigned using canonical epithelial, renal-tumour-lineage, immune and stromal markers within the framework of the source study. In the chRCC specimen, clusters C0-C3 co-expressed EPCAM, KRT7, KRT8, KRT18, RHCG, FOXI1 and ATP6V1B1 with little PTPRC or LYZ signal and were combined as tumour epithelial cells. Clusters C4-C6 were assigned as T/NK cells, tumour-associated macrophages and monocytes. The 69-cell C7 cluster showed mixed epithelial and immune-lineage expression and was excluded from the primary comparisons. Matched normal-kidney clusters 0-2 were combined as renal epithelium and cluster 3 as myeloid.

Tumour epithelial cells in pRCC and ccRCC were identified using epithelial keratins together with subtype-associated markers.

#### GSE159115 specimen assignment and annotation verification

Processed 10x Genomics HDF5 matrices and deposited annotation tables were linked by barcode and sample identifier. Specimen identity and atlas membership were established by cross-referencing GEO records with the source publication's Dataset S1 and confirming barcode-to-annotation concordance. The chRCC specimen was GSM4819732 (SI_21561; donor SS_2016). The deposited cell set and cell-type labels were reproduced without de novo clustering. Total UMI count, detected genes and mitochondrial transcript fraction were recalculated for every library and verified against the deposited values. Normal IC-A cells and ccRCC tumour cells were used as donor-level reference panels; all annotated chRCC compartments were retained for localisation.

#### DUSP15 quantification and donor-level comparisons

DUSP15 abundance was calculated from aggregated raw UMI counts as CPM = DUSP15 UMI/total UMI × $10^6$. Detection prevalence was the proportion of cells with at least one DUSP15 UMI. Donor-level expression was calculated as $log2[(DUSP15UMI + 0.5)/(totalUMI + 1)x10^6]$. Within GSE152938, the chRCC tumour-epithelial compartment was compared with patient-matched normal renal epithelium, pRCC tumour epithelium and the equal-donor mean of two ccRCC tumour-epithelial compartments. Within GSE159115, the chRCC tumour was compared with the equal-donor mean of normal IC-A cells from five donors and ccRCC tumour cells from seven donors. No cell-level P values were used for between-patient inference.

#### Sensitivity and genome-wide contextualisation

The GSE159115 chRCC-versus-normal contrast was assessed using normal IC-A and pooled intercalated-cell references, sequential omission of each normal donor, 1,000 donor-balanced downsampling iterations, restriction to common cellular UMI support and depth-adjusted logistic and quasi-Poisson models. A limma-voom reference-outlier analysis ranked DUSP15 among genome-wide descriptive effects against normal IC-A, pooled normal intercalated-cell and ccRCC tumour reference panels. These rankings characterise the index chRCC specimen and do not constitute a population-level chRCC test.

#### Single-cell immune-metabolic analysis

Raw UMI counts were aggregated by donor and annotated compartment. Gene expression was calculated as $log_2[(count + 0.5)/(totalUMI + 1) \times 10^6]$. Seven Hallmark gene sets from MSigDB release 2025.1 [3] were prespecified from the bulk-RNA-seq results: oxidative phosphorylation, MYC targets V1, MYC targets V2, DNA repair, interferon-γ response, TNF-α signalling via NF-κB and inflammatory response. The first four formed the metabolic/proliferative programme and the final three formed the immune/inflammatory programme.

For each comparison, a gene-level statistic was calculated as the chRCC tumour-epithelial $\log_2$ expression minus the mean $\log_2$ expression of the reference donors. Competitive enrichment was evaluated with cameraPR [4] using ranked statistics. The normalised rank-enrichment score was $2[(meanpathway - generank)/(N+1) - 0.5]$ , where positive values indicate preferentially higher expression in chRCC. Benjamini–Hochberg false-discovery rates were calculated across the seven prespecified pathways within each contrast. Cross-study concordance was summarised with Spearman correlation across the seven rank-enrichment scores.

Counts were normalised to 10,000 UMIs per cell and log-transformed. For each Hallmark set, the mean normalised expression of detected member genes was residualised against log-transformed total UMI count and detected-gene count and then standardised. The four metabolic/proliferative scores and three immune/inflammatory scores were averaged separately to generate composite scores. Continuous associations between log-normalised DUSP15 expression and composite scores were summarised by Spearman correlation before and after the same depth adjustment. Colour ranges in the UMAP displays were restricted to the 2nd–98th percentiles for visualisation only.

**Software and reproducibility**

Analyses were performed in R 4.6.0 using data.table 1.18.4, Matrix 1.7-5, Seurat 5.5.0, SeuratObject 5.4.0, edgeR 4.10.0, limma 3.68.2, rhdf5 2.56.0 and ggplot2 4.0.3. Random seeds were fixed before clustering, UMAP generation and repeated downsampling.

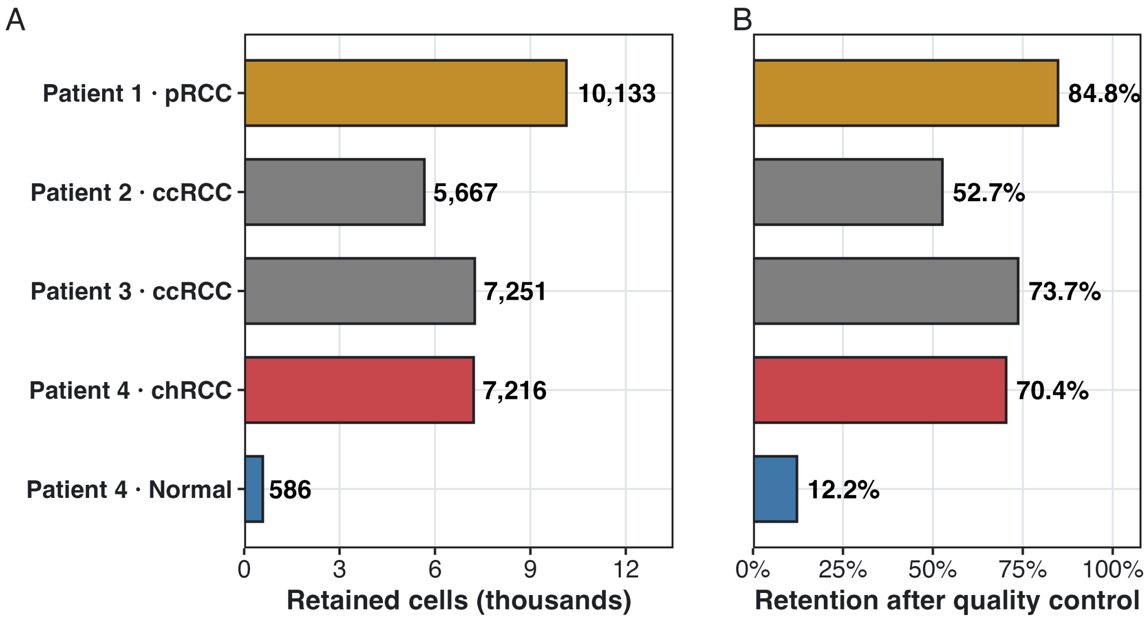


Supplementary Fig. S1. Quality-control retention in the GSE152938 primary dataset

(A) Number of cells retained from each specimen after study-specific quality-control filtering. (B) Percentage of input cell barcodes retained after filtering. Colours denote renal tumour subtype or matched normal kidney.

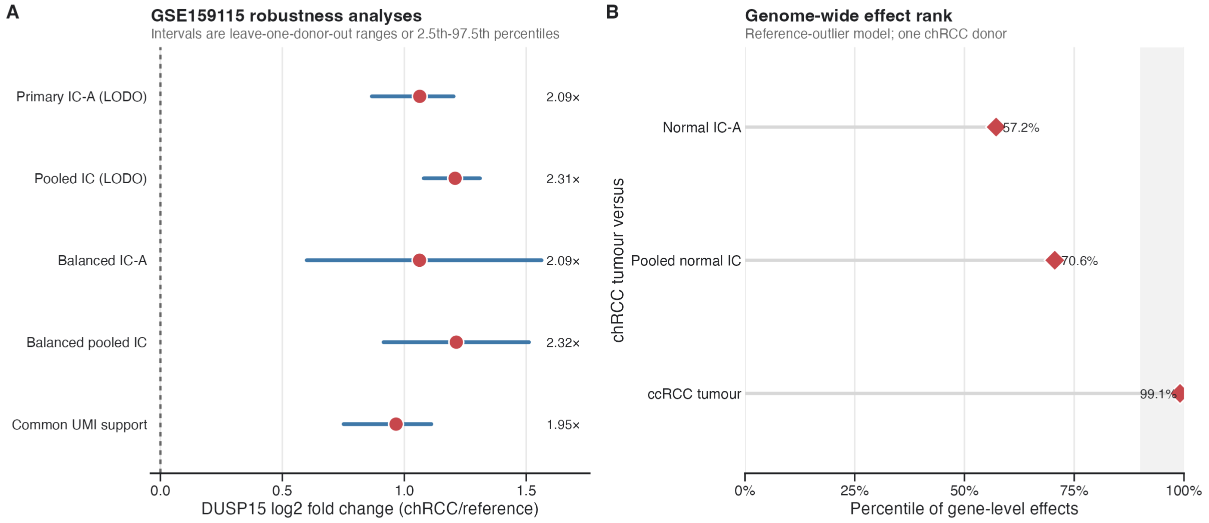


### Supplementary Fig. S2. Sensitivity analyses and genome-wide context in GSE159115

(A) Descriptive DUSP15 $\log_2$ fold changes for the chRCC tumour relative to normal intercalated-cell references across primary, pooled, donor-balanced and common-UMI analyses. Intervals show leave-one-donor-out ranges or the 2.5th–97.5th percentiles from 1,000 downsampling iterations. (B) Percentile rank of DUSP15 among genome-wide descriptive effects for the index chRCC tumour relative to normal IC-A, pooled normal intercalated cells and ccRCC tumour cells. The shaded region denotes the top decile. These are reference-outlier summaries for one chRCC donor.

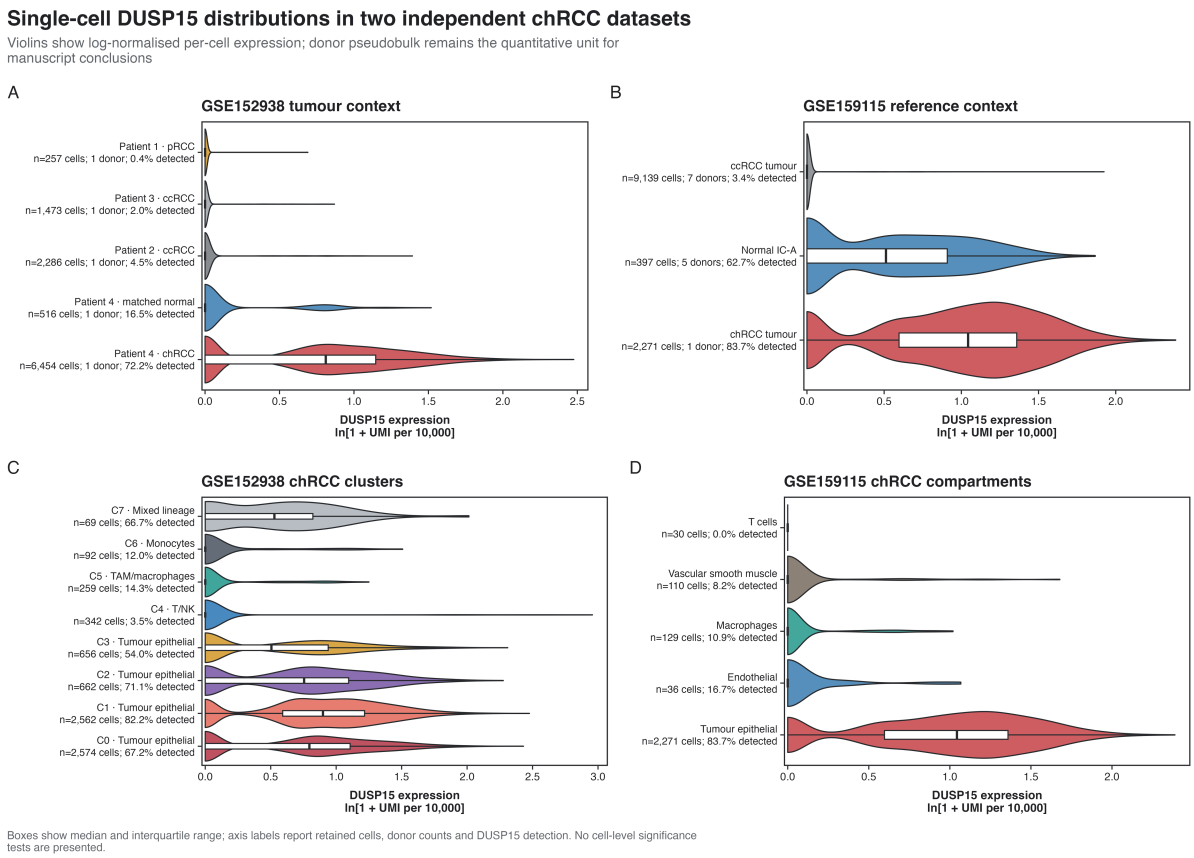


### Supplementary Fig. S3. Single-cell DUSP15 expression distributions in two independent datasets

(A) GSE152938 tumour-epithelial expression distributions across chRCC, matched normal renal epithelium, ccRCC and pRCC. (B) GSE159115 expression distributions in chRCC tumour cells, normal IC-A cells and ccRCC tumour cells. (C) GSE152938 chRCC cluster-level distributions. The mixed-lineage C7 cluster is shown for transparency but was excluded from primary comparisons. (D) GSE159115 chRCC compartment-level distributions. Violin width represents the density of log-normalised per-cell expression, boxes show the median and interquartile range, and labels report cell numbers, donor numbers and DUSP15 detection prevalence. No cell-level significance tests were performed; donor pseudobulk remained the quantitative unit for between-specimen comparisons.

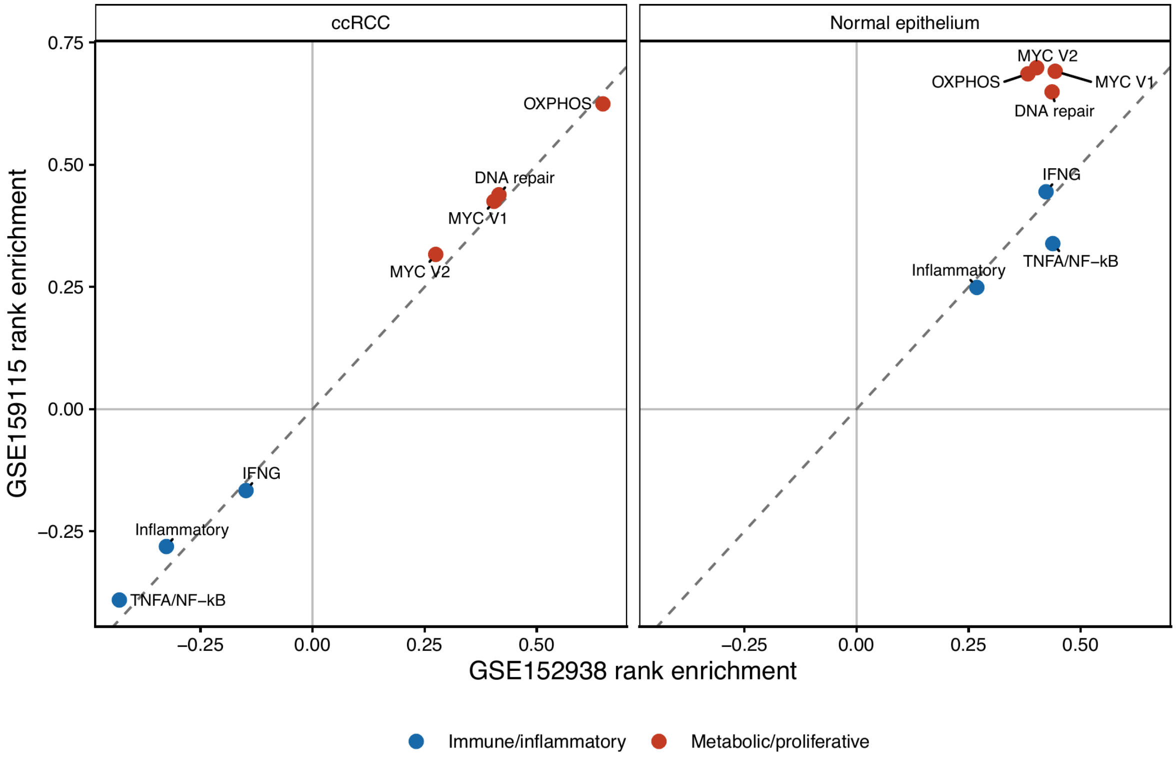


### Supplementary Fig. S4. Cross-study pathway concordance

Rank-enrichment scores from GSE152938 are plotted against the corresponding scores from GSE159115 for chRCC tumour epithelium relative to ccRCC tumour epithelium (left) and normal renal epithelial references (right). The dashed diagonal denotes equality between studies. Red symbols indicate metabolic/proliferative pathways, and blue symbols indicate immune/inflammatory pathways. Spearman correlations were calculated across the seven prespecified Hallmark pathways. Each dataset contained one chRCC donor; the analysis therefore evaluates reproducibility of pathway ordering between the two index specimens rather than estimating a population-level chRCC effect.

### Supplementary Table S9. GSE152938 quality-control and clustering parameters

| GSM | Sample | Patient | Input | Retained | Retention | QC rule | Clustering |
|---|---|---|---|---|---|---|---|
| GSM4630027 | pRCC | Patient 1 | 11949 | 10133 | 84.8% | 200–5,000 genes; mt ≤10% | 20 PCs; res. 0.60 |
| GSM4630028 | ccRCC | Patient 2 | 10762 | 5667 | 52.7% | 200–5,000 genes; mt ≤10% | 25 PCs; res. 0.60 |
| GSM4630029 | ccRCC | Patient 3 | 9837 | 7251 | 73.7% | 200–5,000 genes; mt ≤10% | 25 PCs; res. 0.60 |
| GSM4630030 | chRCC | Patient 4 | 10254 | 7216 | 70.4% | 200–5,000 genes; mt ≤30% | 20 PCs; res. 0.40 |
| GSM4630031 | Normal | Patient 4 | 4800 | 586 | 12.2% | 200–2,500 genes; mt ≤30% | 20 PCs; res. 0.25 |

**mt, mitochondrial transcript percentage; PCs, principal components; res., Louvain resolution.**

### Supplementary Table S10. GSE152938 cluster-to-compartment assignments

| GSM | Clusters | Compartment | Cells | Marker rationale |
|---|---|---|---|---|
| GSM4630027 | 13 | pRCC tumour epithelial | 257 | Epithelial keratins; CDKN2A/SPOCK1/PTGIS |
| GSM4630028 | 1, 2, 5, 11 | ccRCC tumour epithelial | 2,286 | Epithelial keratins; CA9/NDUFA4L2 |
| GSM4630029 | 3, 6, 11 | ccRCC tumour epithelial | 1,473 | Epithelial keratins; CA9/NDUFA4L2 |
| GSM4630030 | 0-3 | chRCC tumour epithelial | 6,454 | EPCAM/KRT7/KRT8/RHCG/FOXI1/ATP6V1B1 |
| GSM4630030 | 4-6 | T/NK; TAM; monocytes | 693 | NKG7; C1QA/LYZ; LYZ |
| GSM4630030 | 7 | Mixed lineage | 69 | Mixed epithelial and immune signal; excluded |
| GSM4630031 | 0-2 | Normal renal epithelial | 516 | EPCAM/KRT8; aggregate renal epithelium |
| GSM4630031 | 3 | Myeloid | 70 | PTPRC/LYZ |

**Unlisted tumour-sample clusters were classified as other tumour microenvironment and excluded from tumour-epithelial subtype comparisons.**

### Supplementary Table S11. DUSP15 by primary compartment in GSE152938

| GSM | Subtype | Compartment | Cells | DUSP15 UMI | CPM | Detected |
|---|---|---|---|---|---|---|
| GSM4630027 | pRCC | Tumour epithelial | 257 | 1 | 0.34 | 0.4% |
| GSM4630028 | ccRCC | Tumour epithelial | 2286 | 107 | 5.45 | 4.5% |
| GSM4630029 | ccRCC | Tumour epithelial | 1473 | 30 | 1.73 | 2.0% |
| GSM4630030 | chRCC | Tumour epithelial | 6454 | 10404 | 144.90 | 72.2% |
| GSM4630030 | chRCC | Monocytes | 92 | 21 | 50.84 | 12.0% |
| GSM4630030 | chRCC | TAM/macrophages | 259 | 74 | 30.88 | 14.3% |
| GSM4630030 | chRCC | T/NK cells | 342 | 12 | 21.43 | 3.5% |
| GSM4630031 | Normal | Normal renal epithelial | 516 | 96 | 22.42 | 16.5% |
| GSM4630031 | Normal | Myeloid | 70 | 0 | 0.00 | 0.0% |

**Values were calculated from aggregated raw UMI counts. The 69-cell mixed-lineage chRCC cluster was excluded.**

### Supplementary Table S12. GSE152938 descriptive chRCC reference effects

| Reference | Reference donors | chRCC $log_2$ CPM | Reference mean | $log_2$ FC | Fold |
|---|---|---|---|---|---|
| Matched normal renal epithelial | 1 | 7.179 | 4.494 | 2.685 | 6.43 |
| pRCC tumour epithelial | 1 | 7.179 | -0.985 | 8.164 | 286.81 |
| ccRCC tumour epithelial | 2 | 7.179 | 1.633 | 5.546 | 46.72 |

**All comparisons contain one chRCC donor and are descriptive.**

### Supplementary Table S13. GSE159115 specimen metadata and atlas assignments

| GSM | Sample | Donor | Atlas | Dataset S1 tissue | GEO tissue | Cross-source discrepancy |
|---|---|---|---|---|---|---|
| GSM4819725 | SI_18854 | SS_2005 | ccRCC | Tumor | Tumor | FALSE |
| GSM4819726 | SI_18856 | SS_2006 | normal | Normal | Tumor | TRUE |
| GSM4819727 | SI_18855 | SS_2006 | ccRCC | Tumor | Benign Adjacent | TRUE |
| GSM4819728 | SI_19704 | SS_2007 | normal | Normal | Tumor | TRUE |
| GSM4819729 | SI_19703 | SS_2007 | ccRCC | Tumor | Benign Adjacent | TRUE |
| GSM4819730 | SI_21255 | SS_2014 | normal | Normal Cortex | Benign Adjacent Cortex | FALSE |
| GSM4819731 | SI_21256 | SS_2014 | normal | Normal Medulla | Benign Adjacent Medulla | FALSE |
| GSM4819732 | SI_21561 | SS_2016 | chRCC | Tumor | Tumor | FALSE |
| GSM4819733 | SI_22369 | SS_2017 | normal | Normal Cortex | Tumor | TRUE |
| GSM4819734 | SI_22368 | SS_2017 | ccRCC | Tumor | Benign Adjacent | TRUE |
| GSM4819735 | SI_22605 | SS_2022 | normal | Normal | Tumor | TRUE |
| GSM4819736 | SI_22604 | SS_2022 | ccRCC | Tumor | Benign Adjacent | TRUE |
| GSM4819737 | SI_23459 | SS_2023 | ccRCC | Tumor | Tumor | FALSE |
| GSM4819738 | SI_23843 | SS_2026 | ccRCC | Tumor | Tumor | FALSE |

Atlas membership was assigned from the source-study supplementary metadata and deposited annotation structure. A cross-source discrepancy denotes a differing individual GEO tissue description; GSM4819732 is the chRCC specimen.

#### Supplementary Table S14. GSE159115 retained cohort summary

| Atlas | Cells | Libraries | Donors | Median UMI | Median genes | Median mt |
|---|---|---|---|---|---|---|
| ccRCC | 20748 | 7 | 7 | 7205.5 | 2158 | 3.3% |
| chRCC | 2580 | 1 | 1 | 13529.5 | 3034 | 16.5% |
| normal | 6146 | 6 | 5 | 4970.0 | 1559 | 6.4% |

**The retained cell set reproduced the deposited annotations after count-metric concordance checking.**

### Supplementary Table S15. DUSP15 by compartment in the GSE159115 chRCC specimen

| Deposited annotation | Cells | DUSP15 UMI | CPM | Cells detected |
|---|---|---|---|---|
| Tumor | 2271 | 6762 | 204.32 | 83.7% |
| Endo | 36 | 12 | 16.44 | 16.7% |
| Macro | 129 | 14 | 12.32 | 10.9% |
| vSMC | 110 | 10 | 9.83 | 8.2% |
| Tcell | 30 | 0 | 0.00 | 0.0% |
| ua | 4 | 8 | 274.73 | 50.0% |

**The four-cell unassigned group is included for completeness and was not used as a biological reference.**

### Supplementary Table S16. GSE159115 donor-level descriptive reference effects

| Reference | $log_2$ FC | Fold | Reference donors | LODO $log_2$ FC | Empirical rank P |
|---|---|---|---|---|---|
| Normal IC-A | 1.063 | 2.09 | 5 | 0.867 to 1.202 | 0.167 |
| Normal pooled IC-A/IC-B/IC-PC | 1.208 | 2.31 | 5 | 1.080 to 1.310 | 0.167 |
| ccRCC tumour | 6.850 | 115.40 | 7 | 6.417 to 7.138 | 0.125 |

**Empirical values are finite-panel reference-outlier ranks and are not population-level chRCC tests. LODO, leave one reference donor out.**

#### Supplementary Table S17. GSE159115 sensitivity analyses

| Analysis | Estimate | Range / naive CI | Interpretation |
|---|---|---|---|
| Balanced IC-A | 1.063 | 0.600 to 1.563 | 1,000 iterations; all positive |
| Balanced pooled IC | 1.213 | 0.915 to 1.511 | 1,000 iterations; all positive |
| Common UMI support | 0.966 | 0.751 to 1.111 | LODO range |
| Depth-adjusted detection | 2.877 | 2.148 to 3.854 | Odds ratio; cell-level diagnostic |
| Depth-adjusted UMI rate | 1.990 | 1.778 to 2.227 | Rate ratio; cell-level diagnostic |

**The first three estimates are $\log_2$ fold changes. Cell-level model intervals are reported only as depth-sensitivity diagnostics and are not donor-valid confidence intervals.**

#### Supplementary Table S18. Cross-study chRCC tumour-epithelial replication

| Dataset | Patient | Tumour epithelial cells | DUSP15 UMI | CPM | Detected |
|---|---|---|---|---|---|
| GSE159115 | SS_2016 | 2271 | 6762 | 204.32 | 83.7% |
| GSE152938 | Patient 4 | 6454 | 10404 | 144.90 | 72.2% |

**The two rows represent independent studies and independent chRCC patients.**

**Supplementary Table S19. Cohorts and reference groups used for immune-metabolic analysis**

| Dataset | Cell population | Donors | Cells | Analytical role |
|---|---|---|---|---|
| GSE152938 | chRCC tumour epithelium | 1 | 6,454 | Index chRCC |
| GSE152938 | Matched normal renal epithelium | 1 | 516 | Normal reference |
| GSE152938 | ccRCC tumour epithelium | 2 | 3,759 | Subtype reference |
| GSE152938 | pRCC tumour epithelium | 1 | 257 | Subtype reference |
| GSE159115 | chRCC tumour epithelium | 1 | 2,271 | Index chRCC |
| GSE159115 | Normal IC-A | 5 | 397 | Lineage reference |
| GSE159115 | ccRCC tumour epithelium | 7 | 9,139 | Subtype reference |

**Each dataset contains one chRCC donor; donor is the biological unit for between-specimen comparisons.**

#### Supplementary Table S20. Prespecified Hallmark gene sets

| MSigDB gene set | Pathway | Programme | Bulk direction | Genes detected |
|---|---|---|---|---|
| HALLMARK_OXIDATIVE_PHOSPHORYLATION | Oxidative phosphorylation | Metabolic/proliferative | Higher | 200 |
| HALLMARK_MYC_TARGETS_V1 | MYC targets V1 | Metabolic/proliferative | Higher | 195 |
| HALLMARK_MYC_TARGETS_V2 | MYC targets V2 | Metabolic/proliferative | Higher | 57 |
| HALLMARK_DNA_REPAIR | DNA repair | Metabolic/proliferative | Higher | 149 |
| HALLMARK_INTERFERON_GAMMA_RESPONSE | IFN-γ response | Immune/inflammatory | Lower | 196 |
| HALLMARK_TNFA_SIGNALING_VIA_NFKB | TNF-α/NF-κB | Immune/inflammatory | Lower | 198 |
| HALLMARK_INFLAMMATORY_RESPONSE | Inflammatory response | Immune/inflammatory | Lower | 200 |

**Bulk direction denotes the direction prespecified from the bulk KICH DUSP15-associated analysis. Gene counts are the detected members used in competitive testing.**

#### Supplementary Table S21a. Competitive pathway rank-enrichment results in GSE152938

| Reference | Pathway | Score | Direction | FDR | Matches bulk |
|---|---|---|---|---|---|
| Matched normal | Oxidative phosphorylation | 0.382 | Up | 2.11e-08 | Yes |
| Matched normal | MYC targets V1 | 0.443 | Up | 3.38e-10 | Yes |
| Matched normal | MYC targets V2 | 0.402 | Up | 1.59e-05 | Yes |
| Matched normal | DNA repair | 0.436 | Up | 1.81e-09 | Yes |
| Matched normal | IFN-γ response | 0.423 | Up | 1.03e-09 | No |
| Matched normal | TNF-α/NF-κB | 0.438 | Up | 3.38e-10 | No |
| Matched normal | Inflammatory response | 0.268 | Up | 7.06e-05 | No |
| ccRCC | Oxidative phosphorylation | 0.648 | Up | 1.73e-20 | Yes |
| ccRCC | MYC targets V1 | 0.405 | Up | 8.53e-09 | Yes |
| ccRCC | MYC targets V2 | 0.275 | Up | 0.004 | Yes |
| ccRCC | DNA repair | 0.416 | Up | 1.54e-08 | Yes |
| ccRCC | IFN-γ response | -0.148 | Down | 0.030 | Yes |
| ccRCC | TNF-α/NF-κB | -0.431 | Down | 1.06e-09 | Yes |
| ccRCC | Inflammatory response | -0.326 | Down | 2.54e-06 | Yes |
| pRCC | Oxidative phosphorylation | 0.751 | Up | 7.01e-28 | Yes |
| pRCC | MYC targets V1 | 0.533 | Up | 1.32e-14 | Yes |
| pRCC | MYC targets V2 | 0.512 | Up | 5.29e-08 | Yes |
| pRCC | DNA repair | 0.478 | Up | 5.19e-11 | Yes |
| pRCC | IFN-γ response | 0.106 | Up | 0.119 | No |
| pRCC | TNF-α/NF-κB | -0.135 | Down | 0.054 | Yes |
| pRCC | Inflammatory response | -0.166 | Down | 0.020 | Yes |

**Positive scores indicate preferentially higher pathway-gene ranks in the index chRCC tumour epithelium. FDR was calculated across the seven prespecified pathways within each contrast. Results describe one chRCC donor per dataset.**

#### Supplementary Table S21b. Competitive pathway rank-enrichment results in GSE159115

| Reference | Pathway | Score | Direction | FDR | Matches bulk |
|---|---|---|---|---|---|
| Normal IC-A | Oxidative phosphorylation | 0.686 | Up | 5.02e-23 | Yes |
| Normal IC-A | MYC targets V1 | 0.691 | Up | 1.86e-23 | Yes |
| Normal IC-A | MYC targets V2 | 0.699 | Up | 6.92e-14 | Yes |
| Normal IC-A | DNA repair | 0.649 | Up | 3.17e-19 | Yes |
| Normal IC-A | IFN-γ response | 0.445 | Up | 7.47e-11 | No |
| Normal IC-A | TNF-α/NF-κB | 0.339 | Up | 6.49e-07 | No |
| Normal IC-A | Inflammatory response | 0.249 | Up | 2.29e-04 | No |
| ccRCC | Oxidative phosphorylation | 0.625 | Up | 2.54e-18 | Yes |
| ccRCC | MYC targets V1 | 0.425 | Up | 2.80e-09 | Yes |
| ccRCC | MYC targets V2 | 0.317 | Up | 8.88e-04 | Yes |
| ccRCC | DNA repair | 0.439 | Up | 4.31e-09 | Yes |
| ccRCC | IFN-γ response | -0.167 | Down | 0.016 | Yes |
| ccRCC | TNF-α/NF-κB | -0.390 | Down | 2.51e-08 | Yes |
| ccRCC | Inflammatory response | -0.281 | Down | 6.20e-05 | Yes |

**Positive scores indicate preferentially higher pathway-gene ranks in the index chRCC tumour epithelium. FDR was calculated across the seven prespecified pathways within each contrast. Results describe one chRCC donor per dataset.**

### Supplementary Table S22. Cross-study pathway reproducibility

| Reference family | Spearman rho | Expected direction | FDR < 0.05 in both | Interpretation |
|---|---|---|---|---|
| Normal epithelium | 0.214 | 4/7 | 7/7 | Metabolic directions reproduced; immune direction reversed |
| ccRCC | 1.000 | 7/7 | 7/7 | Metabolic and immune directions reproduced |

**Correlations were calculated across the seven prespecified Hallmark rank-enrichment scores.**

### Supplementary Table S23. Depth-adjusted composite scores by chRCC cellular compartment

| Dataset | Compartment | Cells | Mean metabolic score | Mean immune score |
|---|---|---|---|---|
| GSE152938 | Tumour epithelial | 6,454 | 0.101 | -0.184 |
| GSE152938 | Tumour microenvironment | 693 | -1.012 | 1.713 |
| GSE159115 | Tumour epithelial | 2,271 | 0.114 | -0.238 |
| GSE159115 | Tumour microenvironment | 305 | -0.859 | 1.781 |

**Scores are standardised residual composites. GSE159115 microenvironment summaries exclude four unassigned cells.**

#### Supplementary Table S24. Within-tumour cell-level associations

| Dataset | Comparison | Cells | Raw rho | Depth-adjusted rho |
|---|---|---|---|---|
| GSE152938 | DUSP15 vs metabolic composite | 6454 | 0.013 | 0.079 |
| GSE152938 | DUSP15 vs immune composite | 6454 | 0.005 | -0.096 |
| GSE159115 | DUSP15 vs metabolic composite | 2271 | 0.001 | 0.025 |
| GSE159115 | DUSP15 vs immune composite | 2271 | -0.040 | -0.135 |

**Spearman correlations are descriptive within one tumour specimen; cells are not independent biological replicates.**
